# On predicting heterogeneity in nanoparticle dosage

**DOI:** 10.1101/2022.05.26.493665

**Authors:** Celia V. Dowling, Paula M. Cevaal, Matthew Faria, Stuart T. Johnston

**Affiliations:** School of Mathematics and Statistics, The University of Melbourne, Australia; Department of Microbiology and Immunology, The University of Melbourne at the Peter Doherty Institute for Infection and Immunity, Australia; Department of Biomedical Engineering, The University of Melbourne, Australia

**Keywords:** Nanoparticles, stochastic modelling, heterogeneity, dosage distribution, cell division, nanoparticle-cell interactions

## Abstract

Nanoparticles are increasingly employed as a vehicle for the targeted delivery of therapeutics to specific cell types. However, much remains to be discovered about the fundamental biology that dictates the interactions between nanoparticles and cells. Accordingly, few nanoparticle-based targeted therapeutics have succeeded in clinical trials. One element that hinders our understanding of nanoparticle-cell interactions is the presence of heterogeneity in nanoparticle dosage data obtained from standard experiments. It is difficult to distinguish between heterogeneity that arises from stochasticity in nanoparticle behaviour, and that which arises from heterogeneity in the cell population. Mathematical investigations have revealed that both sources of heterogeneity contribute meaningfully to the heterogeneity in nanoparticle dosage. However, these investigations have relied on simplified models of nanoparticle internalisation. Here we present a stochastic mathematical model of nanoparticle internalisation that incorporates a suite of relevant biological phenomena such as multistage internalisation, cell division, asymmetric nanoparticle inheritance and nanoparticle saturation. Critically, our model provides information about nanoparticle dosage at an individual cell level. We perform model simulations to examine the influence of specific biological phenomena on the heterogeneity in nanoparticle dosage. Under certain modelling assumptions, we derive analytic approximations of the nanoparticle dosage distribution. We demonstrate that the analytic approximations are accurate, and show that nanoparticle dosage can be described by a Poisson mixture distribution with rate parameters that are a function of Beta-distributed random variables. We discuss the implications of the analytic results with respect to parameter estimation and model identifiability from standard experimental data. Finally, we highlight extensions and directions for future research.

## 1. Introduction

Nanoparticle-based therapeutics are at the forefront of precision medicine, as nanoparticles are able to deliver payloads to specific locations in the body [1, 2]. This technology has seen widespread usage in mRNA vaccines for COVID-19, where lipid nanoparticles coated by polyethylene glycol are used to encapsulate mRNA [1]. These lipid nanoparticles are delivered intramuscularly and result in an immune response following cellular internalisation, endosomal escape and cytosolic mRNA delivery [1, 3]. Nanoparticles have significant potential beyond vaccine delivery, such as for the treatment of malignant tumours [4]. If nanoparticles that carry chemotherapeutics interact preferentially with the tumour cell population, the tumour can be effectively treated while the toxic side effects of chemotherapy are ameliorated [4]. However, we do not yet fully understand how the physicochemical properties of a nanoparticle inform its ability to preferentially interact with a specific cell population [2, 4, 5]. Further, we cannot reliably predict the *in vivo* behaviour of nanoparticles in humans from *in vitro* experiments or animal models [2]. As such, few clinical trials of nanoparticle-based therapeutics have succeeded [6]. Significant work remains to achieve a more complete understanding of nanoparticle-cell interactions and improve the development pipeline of nanoparticle-based therapeutics.

One factor that hinders understanding of the interactions between nanoparticles and cells is heterogeneous experimental data [7, 8, 9, 10, 11, 12]. In standard experiments there is heterogeneity in the number of nanoparticles internalised across a cell population, even though experiments are commonly conducted on clonally-identical cell lines [8, 9, 10, 11, 12]. We will refer to heterogeneity in the number of internalised nanoparticles as the *dosage distribution*. The presence of a dosage distribution has implications about the effective treatment of the entire cell population [13]. As with all therapies, there is a threshold dose that must be internalised by an individual cell for that cell to be considered “treated,” known as the therapeutic dose [14] (which, in turn, may be heterogeneous [15]). It is therefore critical to understand whether any heterogeneity in experimental data arises from heterogeneity in the underlying biology of the cell population, or whether the heterogeneity can be attributed to the stochastic nature of nanoparticle motion [8, 9, 10, 11, 12]. If the dosage distribution arises from differences in cell characteristics, a certain portion of the cell population may not receive a therapeutic dose of nanoparticles [8]. Treatment-resistant cell subpopulations, even if rare, can be a critical factor in determining whether a disease progresses or is cured [16, 17].

Mathematical models are a natural tool to unravel the origins of heterogeneity in experimental data [8, 9, 11]. Previous investigations have highlighted how heterogeneous data from *in vitro* experiments arises from a combination of the stochastic nature of nanoparticle motion and interactions, and the biological heterogeneity inherent to cell populations [8, 9]. Nanoparticle motion is treated as stochastic as nanoparticles are sufficiently small to be influenced by molecular collisions, as characterised by the Stokes-Einstein equation [18]. However, previous investigations of heterogeneous nanoparticle data have relied on simplified descriptions of the interactions between nanoparticles and cells. For example, previous studies[8, 9] have considered a single stage association model of nanoparticle-cell interactions, where nanoparticles become irreversibly associated to cells. Note that *association* refers to nanoparticles that are either *bound* to the cell membrane or are *internalised* by the cell [19, 20]. In this simplified model, the dosage distribution resulting from stochastic nanoparticle motion is shown to be Poisson distributed with a rate parameter that corresponds to the rate that nanoparticles arrive at the cells [8, 9]. This result relies on several simplifying assumptions, such as an absence of both cell proliferation and rate-limited internalisation processes. However, in reality, the process by which nanoparticles are internalised is significantly more complicated, and can involve multiple steps that include nanoparticle-membrane binding, cell receptor recruitment, nanoparticle-receptor binding, membrane deformation, and nanoparticle recycling [5, 21]. Models that include some or all of these steps have been presented previously in a deterministic context (see the reviews of Johnston *et al*. [22] and Åberg [23] and references therein, for example). However, these models have not been presented in a stochastic context and hence do not provide insight about the interplay between specific internalisation processes and unavoidable heterogeneity in dosage. Note that here stochasticity refers to randomness in the amount of time required for a particular component of the nanoparticle internalisation process, such as nanoparticle-receptor binding or membrane deformation, to occur. This definition is consistent with previous investigations into the source of heterogeneity in nanoparticle dosage distributions [8, 9]. Studies have demonstrated that the simplifying assumptions made in investigations of heterogeneous dosage distributions are not always appropriate [24, 25]. For example, it has been observed that nanoparticle association can be a saturating process, where the number of nanoparticles associated to an individual cell plateaus at a finite number [24, 26, 27, 28]. This may imply that the rate of nanoparticle association decreases with the number of previously-associated nanoparticles, and hence the dosage distribution may no longer be Poisson [8]. Other potential explanations include that the plateau arises when nanoparticle association and nanoparticle export, via either cell division or recycling, are balanced [27], or that the cells behave differently upon reaching full confluence [29]. Similarly, the inheritance of nanoparticles during cell division is an asymmetric process [10, 25, 30]. Therefore, if any cell division occurs during the course of an experiment, the assumption of a Poisson dosage distribution is no longer appropriate. Additionally, it is currently unclear how the different components of the nanoparticle internalisation process influence the dosage distribution. Consequently, it is not straightforward to predict how much heterogeneity will be present in measured nanoparticle dosage distributions, even in the absence of any confounding measurement noise or heterogeneity in the underlying cell population. If we cannot tell how much of the heterogeneity in dosage is unavoidable in the best case scenario, it is unlikely that we can determine how much of the heterogeneity in dosage can be attributed to heterogeneity in cell characteristics.

Here we present an agent-based model of nanoparticle-cell interactions that provides information about the number of nanoparticles in each phase of the internalisation process at an individual cell level. The model includes relevant cell behaviour, such as the ability for cells to move and to undergo division via cell cycle progression. We examine the nanoparticle dosage distribution that arises in a range of scenarios, including where multistage internalisation, rate-limited internalisation, and cell division are relevant. We demonstrate that the previous assumption that dosage distributions that arise from stochasticity in nanoparticle internalisation are Poisson distributed is not always appropriate. We present analytic expressions for the expected dosage distributions and the amount of heterogeneity in certain simplified cases. In each case, we verify that the analytic expression describes the behaviour in the agent-based model. Crucially, our work provides a theoretical method for predicting the expected heterogeneity in nanoparticle dosage arising from stochastic internalisation processes. This is distinct from the heterogeneity in dosage that arises from heterogeneity in the characteristics of the underlying cell population. The work here is an important step toward being able to identify any heterogeneity in the underlying cell biology via the comparison of experimental and predicted dosage distributions. Based on the analytic results, we discuss heuristic methods for conducting parameter estimation on experimental data. We highlight the importance of measuring certain experimental quantities to be able to estimate key biological parameters, and to be able to distinguish between biological mechanisms that have a similar influence on nanoparticle internalisation. Finally, we discuss avenues and extensions for future research.

## 2. Methods

A standard experimental approach to examine the therapeutic potential of a novel nanoparticle is a nanoparticle-cell association assay, a schematic of which is presented in Figure 1. [20, 31]. In an association assay, a cell population is seeded in a culture dish and allowed to grow for a specified amount of time [31]. Here we focus on association assays for adherent cell lines, where the cells grow on the bottom of the culture dish, though we note that experiments are also performed for suspension cell lines, where the cells are suspended in culture media. The experiment effectively begins when the cell culture media is replaced with fresh media that contains a suspension of nanoparticles. During the course of the experiment, the nanoparticles move through the fluid and interact with the cells at the bottom of the culture dish [24, 32]. Nanoparticle motion is driven by a combination of mechanisms that can include diffusion, sedimentation, agglomeration and convection [24, 32, 33, 34]. Nanoparticle motion and dosimetry are discussed in detail elsewhere; see the review by Cohen *et al*. [35], for example. If the nanoparticles are sufficiently small and/or light or the culture media is continuously mixed [36], and the nanoparticles are stable in solution, nanoparticle motion can be well-approximated by diffusion. Nanoparticles interact with cells via a range of biological mechanisms, including receptor and surface binding, and can become internalised through a number of endocytic pathways [5, 21]. Techniques can be applied to distinguish between strongly-bound and internalised nanoparticles, though this is not yet standard practice [19]. A common approach for measuring nanoparticle dosage is flow cytometry, which records the fluorescent signal for individual cells [37]. Flow cytometry typically provides information for at least 10^5^ individual cells at each time point in an experiment. The number of nanoparticles per cell can be (approximately) obtained from the fluorescent signal. The output of an association assay is typically the average number of nanoparticles per cell at a number of time points [31]. The output is reported as either bound and internalised nanoparticles, if appropriately distinguished [19], or as associated nanoparticles (i.e. the sum of bound and internalised nanoparticles).

**Figure 1:**
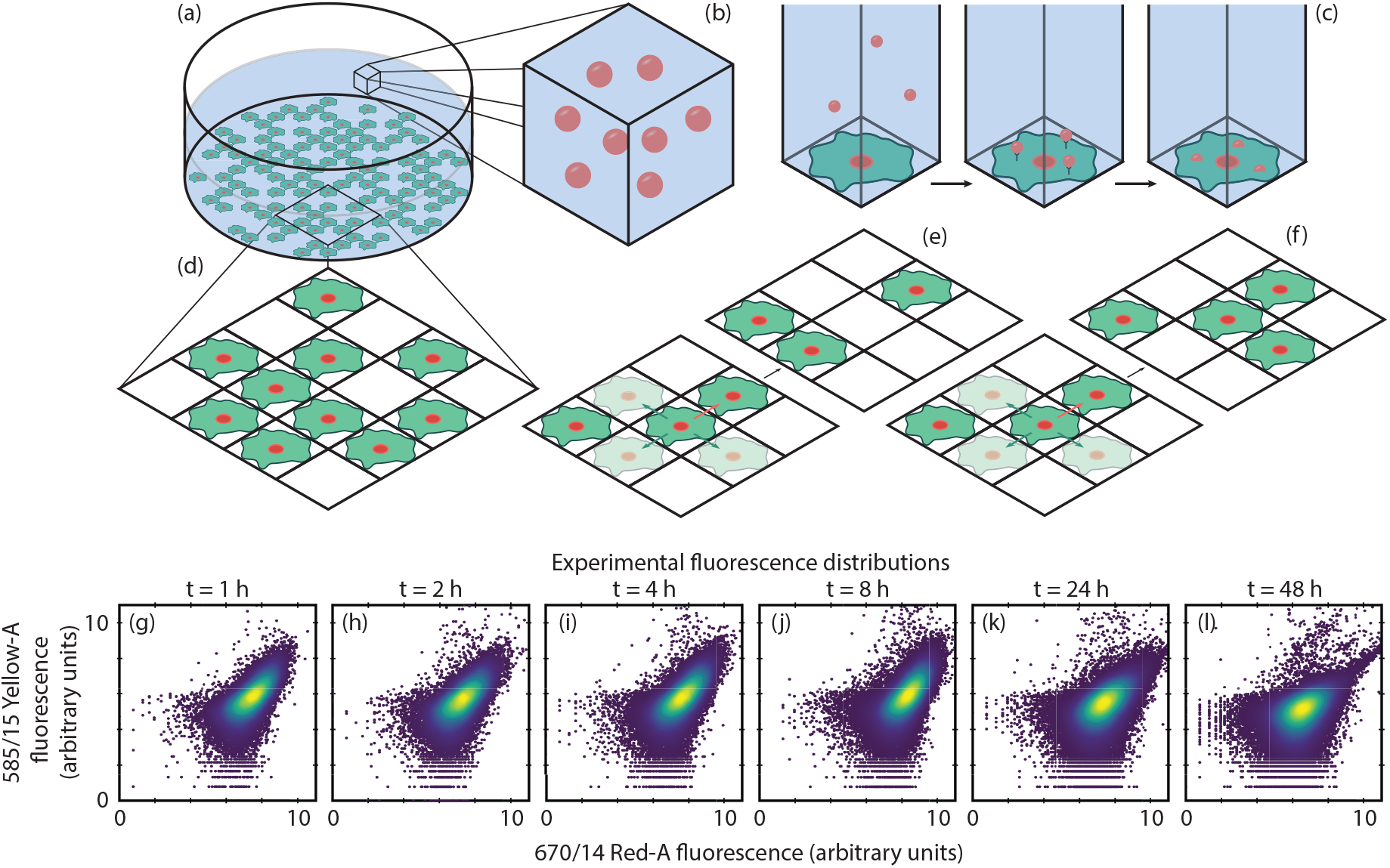
Schematic of standard nanoparticle-cell experiments and the modelling framework. (a) Cells are seeded on a culture dish and (b) immersed in a fluid containing a nanoparticle suspension. (c) Nanoparticles undergo transport through the fluid, bind to the cells, and ultimately become internalised. (d) Cell behaviour on the culture dish is represented via a lattice-based random walk. (e) Example motility events on the lattice, where the cell attempts to move to one of four nearest-neighbour sites. (f) Example proliferation events on the lattice, where the cell attempts to place a daughter cell on one of four nearest-neighbour sites. Green arrows indicate events that will be successful while red arrows indicate events that will not be successful. (g)-(l) Illustrative and representative experimental fluorescence distributions for 100 nm protein G coated iron oxide nanoparticles and Jurkat T cells measured after 1, 2, 4, 8, 24 and 48 hours. Heatmap represents cell frequency (blue - low frequency; yellow - high frequency). The captured fluorescence signals correspond to membrane-bound nanoparticles (585/15 Yellow-A) and associated (i.e. either membrane-bound or internalised) nanoparticles (670/14 Red-A). Experimental details are presented in Appendix A.

Through the use of appropriate mathematical models, parameters that represent the rate of interaction between nanoparticles and cells can be estimated from such data [23, 24, 28]. We present an illustrative example of association assay data, obtained via flow cytometry, in Figures 1(g)-(l). Each data point is the fluorescent signal for an individual cell, which in the appropriate channel is approximately proportional to the number of membrane-bound nanoparticles (585/15 fluorescence axis) and associated nanoparticles (670/14 fluorescence axis) for that cell. Critically, we observe a sizeable spread in the dosage distribution across the cell population. Note that if techniques were not applied to distinguish between membrane-bound and internalised nanoparticles, only the information on the 670/14 fluorescence axis would be available.

Here we develop and introduce a stochastic mathematical model of a nanoparticle-cell association assay. Our modelling framework has two components: a lattice-based random walk for the cell population and a multistage stochastic model of nanoparticle internalisation (Figure 1). Crucially, our modelling framework provides detail at the level of individual cells. This is in contrast to standard modelling approaches, which typically employ ordinary or partial differential equations [22]. Such models are known as population-level models, and provide an estimate of the average number of nanoparticles per cell across the population, rather than detail at an individual cell level. Our framework provides output that is directly comparable with association assay data, both where only association data is available, and where more sophisticated techniques have been employed and hence bound and internalised nanoparticles can be distinguished (Figure 1). Further, the modelling framework captures the stochasticity of nanoparticle internalisation, which is driven by the inherent random motion of the nanoparticles and the stochastic duration of binding and internalisation events [8]. This allows for the prediction of heterogeneity in dosage that arises from inherently stochastic internalisation processes, and is a step toward identifying the heterogeneity in dosage that arises from heterogeneity in cell characteristics.

### 2.1. Cell population model

To describe the behaviour of the adherent cell population, we consider a lattice-based random walk on a two-dimensional square lattice of size *X*Δ by *Y*Δ where Δ is the lattice width [38] (Figure 1(d)). The random walk is chosen to be an exclusion process, where each lattice site may be occupied by, at most, one agent [38, 39]. Here agents represent cells and Δ is chosen to be equivalent to a cell diameter. We impose periodic boundary conditions on the lattice. That is, an agent that leaves the domain is immediately replaced by an identical agent on the opposite side of the domain [40]. The simulation domain can therefore be considered as a representative subsection of a larger experimental domain [40]. It is typically computationally infeasible to simulate the entire experimental domain due to the large number of cells present [40].

Agents on the lattice undergo motility and proliferation events [41]. We make the assumption that cell death does not play a significant role in the experiment, that is, that the nanoparticle dose is not cytotoxic. For a motility event, an agent is randomly selected, with replacement, with probability *M* during a timestep of duration *τ*. The agent, located at (*x*Δ, *y*Δ), attempts to move to a target site that is randomly selected from the four nearest-neighbour sites located at (*x*Δ ± Δ, *y*Δ) and (*x*Δ, *y*Δ ± Δ). The motility event will be successful if the target site is unoccupied, and will otherwise be aborted (Figure 1(e)) [38, 39, 40, 41]. Allowing agents to be motile avoids potential issues with spatial structure that are known to arise in the absence of motility [41], and is consistent with previous assumptions that the nanoparticle concentration in the media does not vary with the *x* and *y* directions [35].

To represent cell division, we consider a multistage proliferation process [42]. Including multiple stages, denoted 1 ≤ *p* ≤ Π, in the proliferation process allows for cell cycle behaviour to be incorporated in the model. The cell cycle is typically considered as four distinct stages: Gap 1 (G1), Synthesis (S), Gap 2 (G2), and Mitosis (M), with cell division occurring at the end of the M phase [43]. Debate continues about whether, and how, the cell cycle influences the rate of nanoparticle internalisation [43, 44, 45]. The simplest model (Π = 1) represents a cell with an exponentially-distributed time between division events. We note that such a simple model is not an accurate reflection of reality, as it possible for a cell in the model to divide multiple times in a short timeframe. However, the assumption of exponential waiting times between events is an oft-overlooked component of translating between stochastic models and ordinary differential equation models [46]. This assumption ensures that the mean behaviour in the simplest form of our stochastic model will be consistent with established models (Appendix B). Moreover, exponential waiting times prove useful in facilitating analytic predictions of dosage distributions. It is, however, important to keep in mind the limitations of the assumption. For Π > 1, that is, multiple proliferation stages, the distribution of cell division times changes from an exponential distribution to a hypoexponential distribution. The distribution of cell division times refers to the time taken between division events for an individual cell. A hypoexponential distribution has been shown to more accurately reflect experimental observations of cell division times [42]. Note that the number of stages in the proliferation model do not necessarily correspond to the four stages in the cell cycle. In our model, an agent at proliferation stage *p* is randomly selected, with replacement, with probability *P_p_* during a timestep of duration *τ*. If *p* ≠ Π, the agent simply progresses to the *p* + 1th stage of the cell cycle [42]. If *p* = Π the agent, located at (*x*Δ, *y*Δ), attempts to place a daughter agent at a target site that is randomly selected from the four nearest-neighbour sites located at (*x*Δ ± Δ, *y*Δ) and (*x*Δ, *y*Δ ± Δ). Similar to motility events, a proliferation event will be successful if the target site is unoccupied, and will otherwise be aborted (Figure 1(f))[41]. If the event is successful, the proliferation stages of both the parent and daughter agent are set to *p* = 1. If the event is aborted, the agent remains in stage Π. Similar models (with Π = 1) have been validated against experimental data of cell migration and proliferation from scratch and barrier assays [40, 47]. A key assumption here is that the cell population exists in a monolayer, that is, that cells do not grow or move over the top of each other. Different modelling assumptions are necessary if the relevant cell population exhibits three-dimensional structure.

### 2.2. Nanoparticle internalisation model

To describe the internalisation of nanoparticles by the cell population, we implement a multistage stochastic process. That is, nanoparticles transition through stages 0 ≤ *s* ≤ *S*, where stage 0 represents nanoparticles in the culture media and stage *S* represents fully internalised nanoparticles. We can select particular *S* values to represent specific models of internalisation. For example, *S* = 1 represents a single stage association model, whereas *S* = 2 represents a model of nanoparticle binding and internalisation. We initially impose a nanoparticle suspension that contains *N*_initial_ nanoparticles. We assume that these nanoparticles are uniformly distributed throughout the experimental domain, that is, that the suspension in the culture media is well-mixed. This assumption is appropriate provided that the media is continuously stirred or the nanoparticles are sufficiently small and/or light such that diffusion is the dominant component of nanoparticle motion [36]. Further, we assume that there are no dominant convective currents in the media and that nanoparticle agglomeration is minimal [48]. However, we note that this is not an appropriate assumption for all nanoparticle systems. Under these assumptions, in the model there are initially, on average, *N*_initial_/*XY* nanoparticles in the media above each lattice site (Figure 1(c)). This implies that nanoparticle diffusion is rapid compared to nanoparticle uptake and hence nanoparticle concentration does not vary in the horizontal directions, consistent with previous dosimetry studies [35, 48].

During a timestep of duration *τ*, an individual nanoparticle in the media can bind to an agent with probability *P*_bind_. This binding probability represents two processes. First, the nanoparticle must reach the cell layer during that timestep via random motion and an agent must be present on the relevant lattice site. Nanoparticle diffusivity arises from the Stokes-Einstein equation [34], and is given by

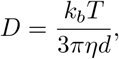

where *k_b_* is the Boltzmann constant, *T* is the temperature of the media, *η* is the dynamic viscosity of the media and *d* is the nanoparticle diameter. The inverse relationship between nanoparticle diffusivity and diameter implies that smaller nanoparticles are more likely to arrive at the cell layer during a timestep. The second process is the nanoparticle binding to the cell, given that the nanoparticle has arrived at the cell layer. This occurs at a rate proportional to the strength of interaction, also known as the affinity, between the nanoparticle and the cell [24]. If there is no cell at the relevant lattice site, the nanoparticle remains in the fluid. The probability that the nanoparticle arrives at the cell layer and binds to a cell during a timestep is a function of media depth, *h*, and nanoparticle diffusivity [49],

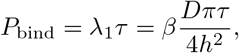

where 0 ≤ *β* ≤ 1 is the probability that the nanoparticle binds to the cell, given it has arrived at the cell layer, and *λ*_1_ is the rate parameter describing the transition from a nanoparticle in the fluid (*s* = 0) to a nanoparticle bound to a cell membrane (*s* = 1). Calculating this probability allows us to determine the rate of nanoparticle binding without explicitly tracking the locations of nanoparticles in the media. We note that this probability may not be appropriate if the relevant nanoparticles exhibit nontrivial amounts of sedimentation or agglomeration.

The first stage of the internalisation process corresponds to nanoparticles binding to either the cell membrane or a receptor. Depending on the physicochemical properties of the nanoparticles and the cell phenotype, the nanoparticle can be internalised via a number of endocytic pathways. These pathways can involve macropinocytosis, receptor recruitment, formation of caveolae or membrane invaginations, and endosomal trafficking [5, 21]. Additionally, the pathway is not necessarily unidirectional, as nanoparticle-receptor binding is reversible and nanoparticle recycling (exocytosis) can occur. As, in general, these pathways are not fully characterised, we choose to represent internalisation as an *S*-stage stochastic process. We make the assumption that each stage of internalisation is a Poisson process, that is, progressing through the stage requires an exponentially-distributed amount of time. We denote the rate parameter of the *s*th stage as *λ_s_*, which represents the rate of transition from stage *s* − 1 to stage *s*. We model the reverse pathway (i.e. nanoparticle-receptor unbinding or nanoparticle exocytosis) by allowing nanoparticles to transition from the *s*th stage of internalisation back to the fluid stage (*s* = 0) with rate parameters 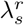.

It has been demonstrated that certain cell lines have a finite carrying capacity for associated nanoparticles [24, 26, 28], though we note that this is not universally observed. It is not always clear whether this bottleneck occurs due to a saturation of nanoparticles at the cell surface, or at the latter stages of internalisation. We account for this phenomenon by incorporating a maximum number of nanoparticles (i.e. a carrying capacity) at each stage of internalisation, *K_s_*, and scaling the corresponding rate parameter by the available fraction of a cell’s carrying capacity, giving an effective rate parameter of 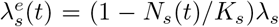, where *N_s_*(*t*) is the number of nanoparticles in the *s*th stage of internalisation at time *t*. The effective rate parameter reduces to the original rate parameter in the limit *K_s_* → ∞, which reflects the case where no saturation occurs in an experiment, or where saturation occurs due to a different mechanism. It is possible that observations of nanoparticle saturation can be attributed to nanoparticle recycling, degradation or cell division [27].

The model provides information about the number of nanoparticles in each internalisation stage for each agent in the population. For the *i*th agent in the population, we denote the number of nanoparticles in each stage as 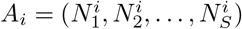. The number of associated nanoparticles for the *i*th agent is the sum of the entries in *A_i_*. Nanoparticles are defined as internalised if they are in stage *S*. When an agent undergoes division, the nanoparticles in each internalisation stage for that agent are split between the parent and daughter agent according to a binomial process. Experimental evidence suggests that a daughter cell inherits between 52% and 72% of the associated nanoparticles [10, 25]. The remainder of the nanoparticles are associated with the parent cell such that the total number of associated nanoparticles is conserved. Accordingly, we randomly distribute the nanoparticles between the daughter and parent agent in the *s*th stage via a binomial process with 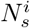 trials and a probability of inheritance *I*. The daughter agent is always defined as the agent that is placed at a neighbouring lattice site.

### 2.3. Model implementation

We perform all simulations of the model on a lattice with *X* = 10 and *Y* = 10 sites in the *x* and *y* directions, respectively. Initially, *C*_0_ unique sites are selected at random and an agent is placed on each site. The cell cycle stage of each agent is selected uniformly at random. We use a fixed timestep approximation to simulate the evolution of the system in time [50]. In each timestep of duration *τ* we update the system according to three types of events: nanoparticle internalisation events, cell motility events, and cell proliferation events. For internalisation events, we keep track of the number of nanoparticles in each stage of internalisation for each agent in the simulation. We assume that nanoparticles are uniformly distributed throughout the culture media and hence, on average,

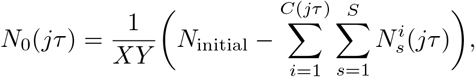

are available in the media above each lattice site after *j* ∈ {ℕ_0_ | *j* ≤ *t*_end_/*τ*} time steps, where *C*(*jτ*) is the number of agents occupying the lattice at the beginning of the *j*th timestep and *t*_end_ is the final experimental time point. This assumption requires that there is no spatial structure in the cell population that could induce a non-uniform distribution of nanoparticles in the *x* and *y* directions; a common feature in nanoparticle dosimetry models [35, 48]. We can satisfy this assumption easily provided that cells are motile. We note that this assumption may not be appropriate for immotile cell lines or for cell lines that form three-dimensional structures. For each nanoparticle in the *s* − 1th stage of internalisation, progression to the next stage occurs with probability 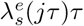. We select *τ* such that *λ_s_τ* ≪ 1 (and *M* ≤ 1, *P_p_* ≤ 1 ∀ *p*), so that it is unlikely that a nanoparticle will progress through more than one internalisation stage in a single timestep. We calculate the number of nanoparticles that progress between internalisation stages for all agents and stages, and then simultaneously update the state of the system. For motility events, *C*(*jτ*) agents are selected at random, with replacement, and attempt to undergo a motility event with probability *M*. After the motility events, an additional *C*(*jτ*) agents are selected at random, with replacement, to proceed to the next stage of the cell cycle or attempt to divide, as appropriate, with probability *P_p_*, where *p* is the cell cycle stage of the selected agent. If the agent divides, the associated nanoparticles are divided between the parent and daughter agent according to the aforementioned binomial process. Recall that both cell motility and division events are only successful if the randomly-selected target site is unoccupied. A number of identically-prepared realisations of the model are performed to generate representative average behaviour; the number of realisations performed to obtain a specific result is presented in the relevant figure caption. All simulations are performed in Matlab R2020b and all code is available on GitHub. For all simulations we use *M* = 1, *τ* = 1/6 h, *t*_end_ = 24 h, *T* = 310 K, *η* = 1.0005 × 10^−3^ kg m^−1^ s^−1^, *k_b_* = 1.38 × 10^−23^ m^2^ kg s^−2^ K^−1^ and Δ = 25 *μ*m. Note that in the code *M* and *P_p_* are calculated by defining an expected waiting time between motility and proliferation events, respectively.

## 3. Results and Discussion

We are interested in the distribution of *N_s_*(*t*) across the agent population, as this represents the dosage distribution for each stage of the internalisation process. It is not always possible to obtain such information experimentally, as it can be difficult to reliably identify which stage of internalisation a nanoparticle is in [19]. However, as we will demonstrate, it is possible to obtain analytic results for the dosage distribution for each stage of internalisation under certain simplifying assumptions. It is important to recognise that the accuracy of the conclusions drawn from the results is contingent on the accuracy of the model assumptions. While we present simulation results, the main focus of this manuscript is to present analytic solutions (i.e. solutions that can be expressed via an equation). Such solutions are more efficient to obtain, and provide greater insight about the direct relationship between the expected heterogeneity and the biological parameters, conditions, and mechanisms. However, obtaining analytic solutions (typically) require additional assumptions, relative to simulations; in each case, we discuss the limitations and implications of the relevant model assumptions.

### 3.1. Nanoparticle association

The simplest form of the model that we consider is an irreversible single stage internalisation process for a cell population that is fully confluent. Here we will use the terminology of *associated* nanoparticles to highlight that there is no distinction between nanoparticles bound to the cell membrane and nanoparticles that have been internalised by the cell [24]. While this is an oversimplification of the internalisation process, it aligns with standard experimental practice [19]. As the cell population is fully confluent, no division events are possible, and nanoparticle inheritance can be neglected. This is not necessarily representative of experimental conditions, which are typically performed below full confluence to avoid changes in cell behaviour [29]. However, we build on the results obtained from this simplifying assumption to consider the more relevant case away from full confluence. We further assume that the impact of the carrying capacity can be neglected, that is, that *K*_1_ → ∞ and hence 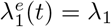. Under these assumptions, the mean number of associated nanoparticles per cell by time *t*

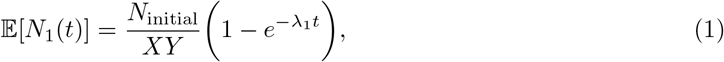

which is the cumulative distribution function (CDF) of an exponential distribution with rate parameter *λ*_1_, multiplied by the average number of nanoparticles initially present in the culture media above each lattice site. As this is simply an exponential process, the distribution of the number of associated nanoparticles per cell by time *t* is a Poisson distribution with rate parameter 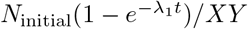, a result that is consistent with previous studies [8, 9]. We present the mean and standard deviation of the number of associated nanoparticles per cell, obtained via simulation, alongside the mean and standard deviation of the Poisson distribution in Figure 2(a). We observe that the analytic predictions of the dosage distribution from Equation (1) matches that obtained from the simulation, both in terms of the mean and standard deviation of the number of associated nanoparticles per cell. The number of associated nanoparticles per cell increases approximately linearly at early time, and plateaus as the number of nanoparticles remaining in the media decreases. While *complete* depletion of the nanoparticles in the media is not a common experimental feature, a sizeable proportion of the initial dosage may associate by the end of the experiment [51], which will influence the effective association rate. As such, it is important to include depletion effects in the model. Dosage depletion can be effectively neglected in the model by selecting a sufficiently large initial dose alongside a sufficiently small rate parameter for association such that only a small proportion of the initial dose of nanoparticles associate over the course of the experiment.

It is desirable to commence experiments before the cell population becomes fully confluent either to observe the influence of the cell cycle and division [43] or to avoid changes in cell metabolism that can occur when a cell population is fully confluent [29]. If the cell population is not fully confluent, then the total number of associated nanoparticles is necessarily reduced (changes in metabolism notwithstanding) as it is possible for nanoparticles to arrive at the base of the culture dish without interacting with a cell. If we assume that the amount of cell proliferation over the course of the experiment is negligible (i.e. that the duration of the experiment is well below the doubling time of the cell line), then the number of associated nanoparticles per cell at time *t* is Poisson distributed with rate parameter

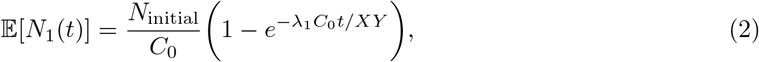

where 0 ≤ *C*_0_ ≤ *XY* is the number of cells in the population. This can be considered as the CDF of an exponential random variable that represents the time taken for a nanoparticle to arrive at the bottom of the culture dish at a location where a cell resides, and to bind to that cell, multiplied by the number of initially available nanoparticles per cell. The rate parameter of the random variable, *λ*_1_*C*_0_/*XY*, decreases with the level of confluence, reflecting that it is less likely for an individual nanoparticle to interact with a cell. While the rate parameter decreases, there is a corresponding increase in the maximum number of nanoparticles per cell (i.e. the number of nanoparticles in the initial suspension is constant but there are fewer cells). Interestingly, the combination of these two effects is that the mean number of associated nanoparticles per cell is approximately independent of confluence when only a small proportion of the nanoparticles are associated (i.e. when *λ*_1_*C*_0_*t/XY* is small and we are in the linear association regime). We recover Equation (1) when *C*_0_ = *XY*.

**Figure 2:**
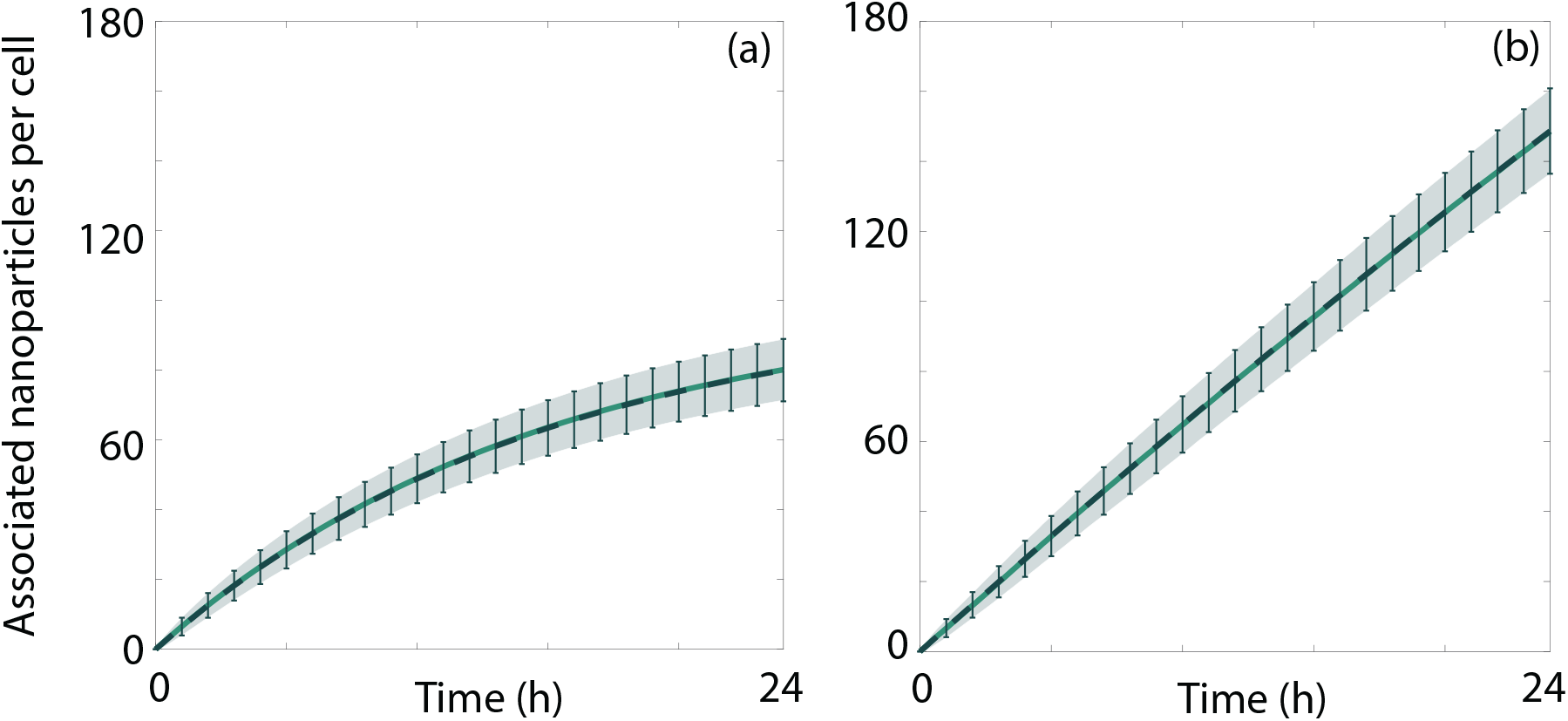
Comparison between the analytic and simulation predictions of the number of associated nanoparticles per cell at (a) full confluence and (b) 10% confluence. Lines correspond to the analytic (dashed) and simulation (solid) predictions of the mean. The analytic and simulation predictions of the mean ± one standard deviation correspond to the error bars and ribbons, respectively. Parameters used are *d* = 20 nm, *N*_initial_ = 10^4^, *h* = 5 × 10^−4^ m, *λ*_1_ = 6.71 × 10^−2^ h^−1^, 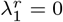, *P_p_* = 0 ∀ *p*, *S* = 1, *K*_1_ → ∞, and (a) *C*_0_ = 100, (b) *C*_0_ = 10. All simulation data are obtained from 200 identically-prepared realisations of the simulation.

We present a comparison between the average simulation behaviour and the Poisson distribution for a cell population at 10% confluence in Figure 2(b). Again, we see that the simulated dosage distribution matches both the analytic mean and standard deviation of the number of associated nanoparticles per cell. Intuitively, the approximately linear behaviour continues for a significantly longer time compared to the fully confluent case as dosage depletion effects are not relevant over the 24 hours of the experiment, due to the reduced number of cells available.

If the cell population is not fully confluent initially, and cell proliferation cannot be neglected, it is not straightforward to obtain a closed-form analytic solution for the mean number of associated nanoparticles per cell. For a single stage model of cell proliferation (i.e. Π = 1), the mean number of cells in the population, 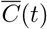, is well-approximated by logistic growth and hence

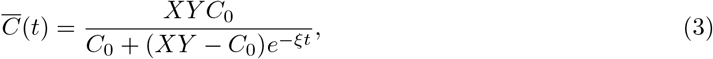

where *ξ* = *P*_1_/*τ* is the rate of cell proliferation. The approximation is accurate provided that the rate of cell motility is significantly higher than the rate of cell proliferation [41]. Certain cell lines are immotile and hence this approximation is not appropriate for all cell lines. To the best of our knowledge, there is no analytic expression for the distribution of the number of cells in the population at time *t* under logistic growth. We can, however, use Equation (3) to find an integral expression for the mean number of associated nanoparticles per cell in the presence of cell proliferation

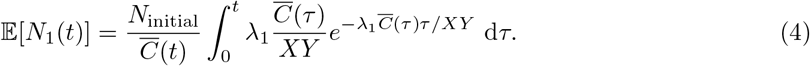

Similar to Eqauation (2), this can be considered as the CDF of a random variable that represents the time required for a nanoparticle to arrive at, and bind to, a cell, multiplied by the number of initially available nanoparticles per cell. However, as the number of cells increases with time, we must account for the increased chance that a nanoparticle arrives at the base of the culture dish and interacts with a cell. In general, this expression cannot be evaluated analytically, and we are restricted to numerical integration methods.

We present a comparison between the mean number of associated nanoparticles per cell, as predicted by the simulation and obtained from numerically integrating Equation (4) for a cell population undergoing proliferation in Figures 3(a) and (c). We observe that the mean number of nanoparticles per cell decreases, compared to the case without proliferation. This result is expected, as the nanoparticles associated to a cell are split between the parent and daughter cell upon division. Further, we observe that the integral expression in Equation (4) accurately describes the mean number of nanoparticles per cell obtained from the simulation. In Figures 3(a) and (c) we see that the ratio of the standard deviation to the mean of the dosage distribution when proliferation is present does not change appreciably when we substantially increase the initial dosage. In contrast, for the case without proliferation, the ratio is significantly decreased for higher initial dosages. This is because, in the absence of proliferation, the dosage distribution has a standard deviation that scales with the square root of the mean dosage, as noted above. As such, we expect to see a (relatively) narrower dosage distribution for higher initial dosages when cell proliferation is not present.

**Figure 3:**
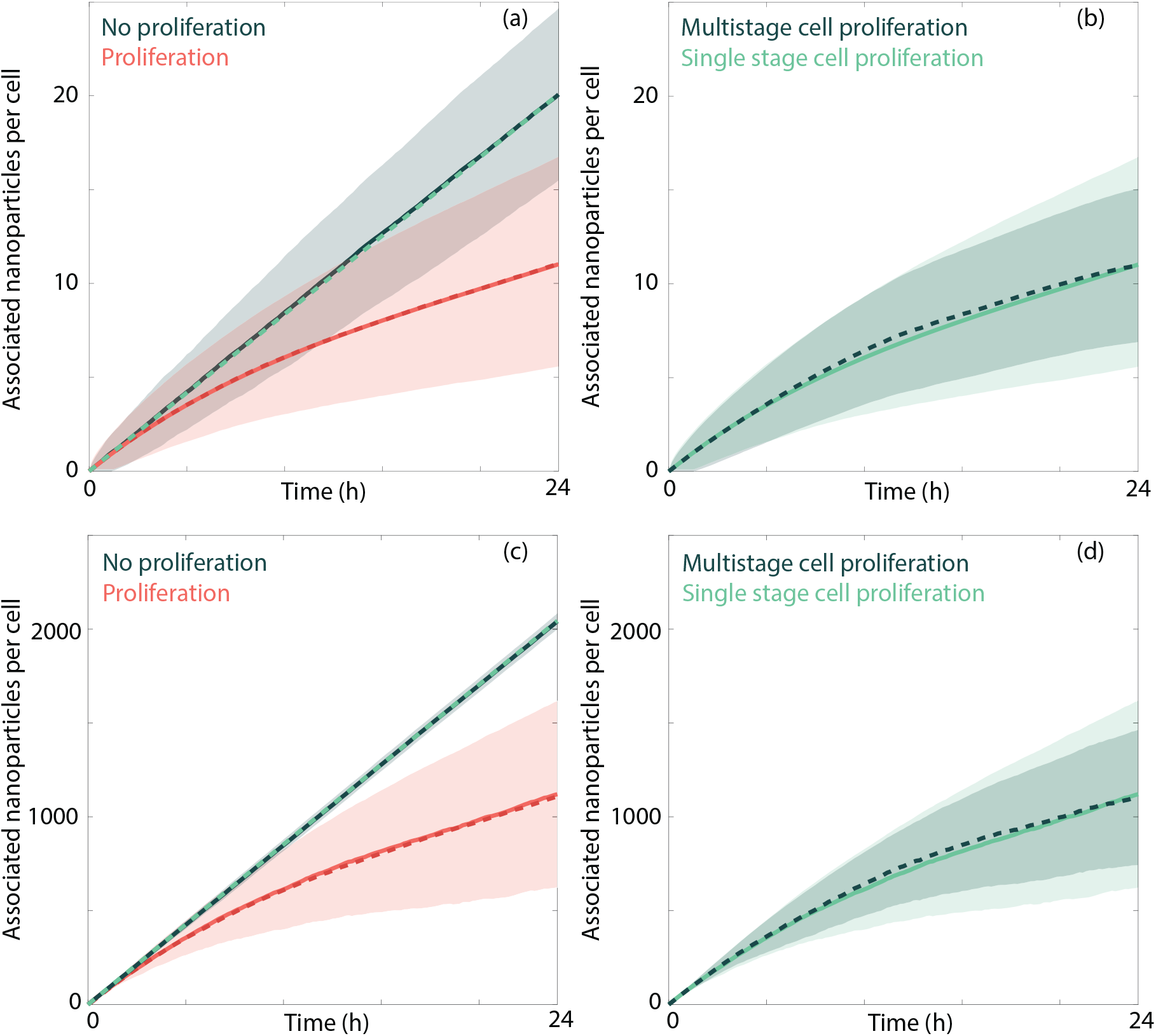
Comparison of the number of associated nanoparticles per cell for different proliferation models. (a),(c) Comparison between no proliferation (green) and single stage proliferation (red) for analytic (dashed) and simulation (solid) predictions. (b),(d) Comparison between single stage (light green, solid) and multistage proliferation (dark green, dashed) for simulation predictions. The ribbons correspond to the mean ± one standard deviation. Parameters used are *d* = 50 nm, *S* = 1, *C*_0_ = 10, *I* = 0.7, *K*_1_ → ∞, 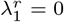, (a)-(b)*N*_initial_ = 10^5^, *h* = 5 × 10^−3^ m, *λ*_1_ = 8.42 × 10^−4^ h^−1^; (c)-(d) *N*_initial_ = 5 × 10^6^, *h* = 3 × 10^−3^ m, *λ*_1_ = 1.70 × 10^−3^ h^−1^; (a),(c): Π = 1, *P*_1_ = 0 (green), *P*_1_ = 1/72 (red); (b),(d): Π = 1 (light green), *P*_1_ = 1/72 (light green), Π = 24 (dark green), *P_p_* = 1/3 ∀ *p* (dark green). All simulation data are obtained from 1000 identically-prepared realisations of the simulation.

The choice of proliferation model has a small influence on the mean number of associated nanoparticles per cell, as can be seen in Figures 3(b) and (d), where we compare simulation results obtained using either a single stage or multistage (Π = 24) model of proliferation. Instead, the influence of the proliferation model manifests in the spread of the dosage distribution. The multistage proliferation model results in a narrower dosage distribution, as the number of divisions per cell is more consistent across the population. As discussed, the multistage proliferation model is more biologically plausible than the single stage proliferation model, but there is a trade-off to be made between the level of realism and the ability to make analytic progress. We note that the level of asymmetry present in the inheritance process of associated nanoparticles does not affect the mean number of nanoparticles per cell, as the rate of association is independent of the number of previously-associated nanoparticles. However, the effect of the inheritance is observable in the dosage distributions, presented in Figure 4. The distribution of associated nanoparticles per cell is noticeably broader if the inheritance is asymmetric (Figures 4(g)-(l) and 4(s)-(x)). In the asymmetric case, we observe that there is a substantial increase in the number of cells that have zero nanoparticles associated by 24 hours, which only occurs rarely for symmetric inheritance. We again observe that the dosage distribution is narrower for the multistage proliferation model (Figures 4(m)-(x)) than the single stage proliferation model (Figures 4(a)-(l)).

**Figure 4:**
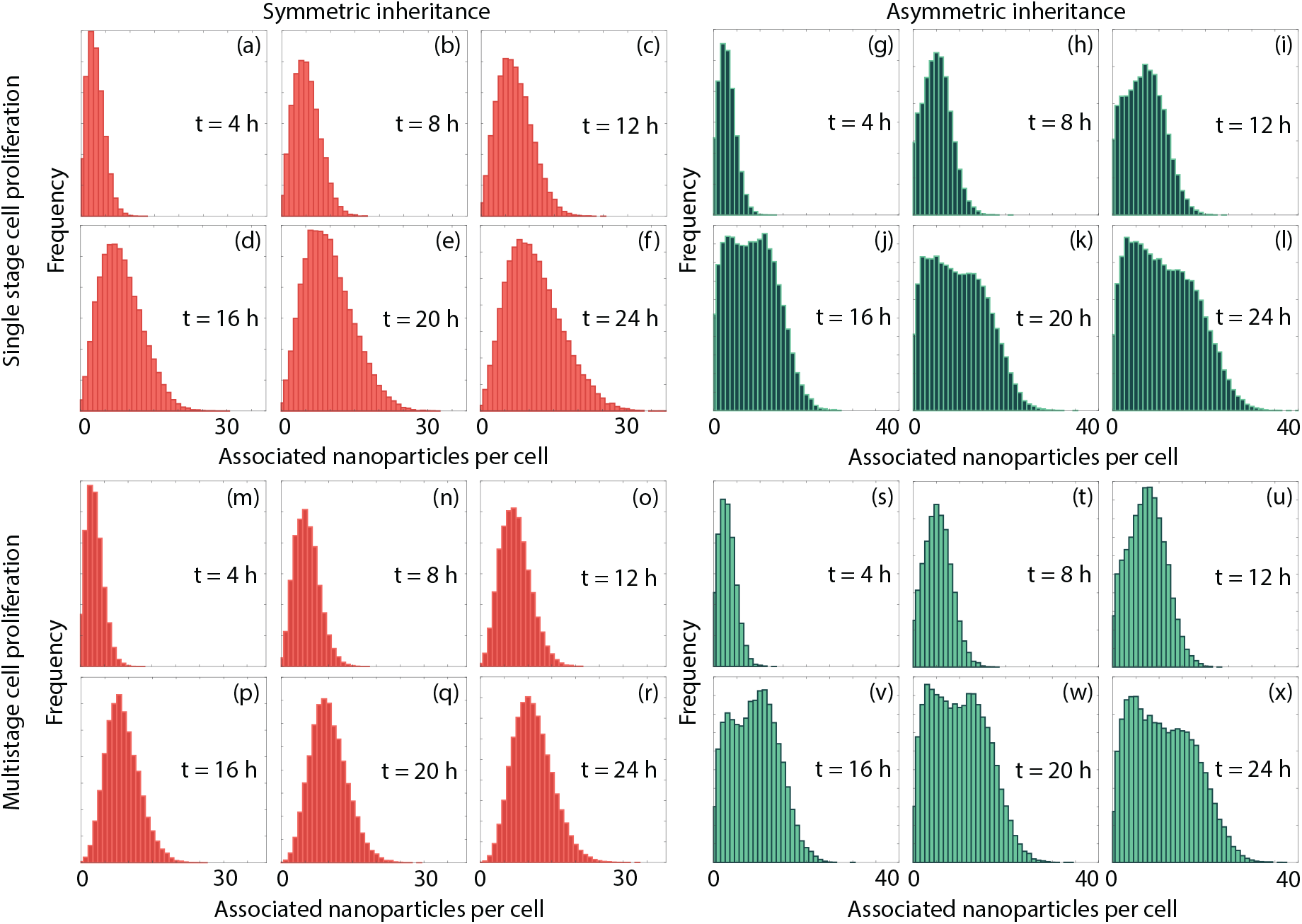
Dosage distributions recorded at four hour intervals for (a)-(f) single stage proliferation and symmetric inheritance, (g)-(l) single stage proliferation and asymmetric inheritance, (m)-(r) multistage proliferation and symmetric inheritance, and (s)-(x) multistage proliferation and asymmetric inheritance. Parameters used are *d* = 50 nm, *N*_initial_ = 10^5^, *h* = 5 × 10^−3^ m, *λ*_1_ = 8.42 × 10^−4^ h^−1^, 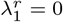, *S* = 1, *C*_0_ = 10, *K*_1_ → ∞, (a)-(f), (m)-(r) *I* = 0.5, (g)-(l), (s)-(x) *I* = 0.9, (a)-(l): Π = 1, *P*_1_ = 1/72; (m)-(x): Π = 24, *P_p_* = 1/3 ∀ *p*. All simulation data are obtained from 1000 identically-prepared realisations of the simulation.

One benefit of employing the model is that it is possible to divide the population dosage distribution into separate dosage distributions based on the number of divisions a cell has experienced. That is, the population dosage distribution can be considered as a mixture distribution of the dosage distributions for cell subpopulations that have never divided, divided once, divided twice, and so on. Here we make the assumption that the cell population is not close to confluence, and hence the growth of the cell population is approximately exponential. Further, we make the assumption that dosage depletion effects can be neglected, that is, that we are in the regime where association is independent of confluence and linear in the absence of proliferation. We first consider the subpopulation of cells that have experienced division exactly once by time *t*, which includes both the parent and daughter cells. We define the time at which the cell undergoes division as *t*_1_. Under the assumption of exponential growth, we know that *t*_1_ is uniformly distributed on the finite interval 0 ≤ *t*_1_ ≤ *t*. This can be seen by defining two independent and identically distributed exponential random variables *X*_1_ and *X*_2_, which represent the time of the first and second divisions of a cell, and calculating *p*(*X*_1_ = *t*_1_|*X*_1_ < *t, X*_1_ + *X*_2_ > *t*) for 0 ≤ *t*_1_ ≤ *t*. Accordingly, the dosage distribution of this subpopulation is a mixture distribution with two equally-weighted components: the dosage distribution of parent cells, which are Poisson distributed with rate parameter

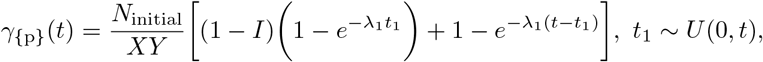

and the dosage distribution of the daughter cells, which are Poisson distributed with rate parameter

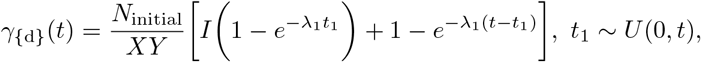

recalling that *I* denotes the proportion of nanoparticles that are inherited by a daughter cell upon division. There are two components for each rate parameter: the number of nanoparticles that associated to the cell after division occurs, and the number of nanoparticles that associated before cell division, and remained after division. There will be, on average, an equal number of daughter and parent cells that have divided exactly once.

We follow a similar process to obtain the dosage distribution for the subpopulation of cells that have divided exactly twice by time *t*. We define the time at which the first and second divisions occur as *t*_1_ and *t*_2_. From order statistics [52], we know that the PDF of *t*_1_ is

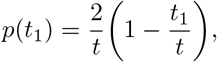

which implies that

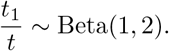

In turn, the time between the first and second division events, *t*_2_ − *t*_1_, is uniformly distributed on 0 ≤ *t*_2_ − *t*_1_ ≤ *t* − *t*_1_. There are four equally-weighted components in the subpopulation dosage mixture distribution, which are Poisson distributed with rate parameters

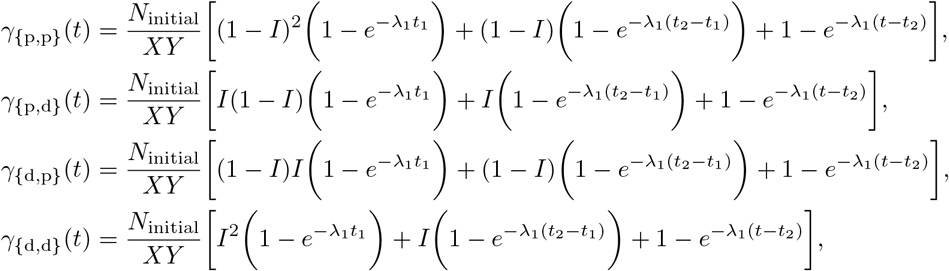

where

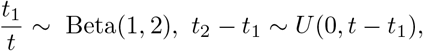

and the subscript of *γ*(*t*) indicates the order and type of division events experienced. For example the subscript {p,d} indicates that a cell has experienced two division events; the first of which where it was the parent cell and the second of which where it was the daughter cell. An increase in the asymmetry of inheritance corresponds to a larger difference between the rate parameters of each component of the mixture distribution, which can induce multimodality in the dosage distribution, as observed in Figure 4. If inheritance is symmetric (i.e. *I* = 0.5), the rate parameters for each component are identical and the dosage distribution is unimodal.

We can generalise this process to obtain the dosage distribution for the subpopulation of cells that have divided *N* times by time *t*, where the time of the *n*th division (1 ≤ *n* ≤ *N*) is denoted *t_n_*. Again, from order statistics [52], the PDF for *t_n_* − *t*_*n*−1_ (i.e. the time between division events) is

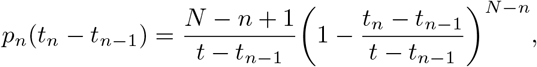

noting that we define *t*_0_ = 0, and hence

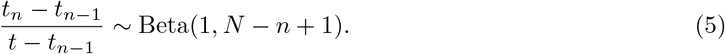

To obtain the general form of the rate parameters, we define the sequence *a_n_* = {*b*_1_, *b*_2_, · · ·, *b_n_*}, where *b_j_* ∈ {d,p}, which represents the order of the *n* division events that a cell has experienced, and whether it was the daughter (d) or parent (p) cell at each division. We define Ω_d,*i*_ and Ω_p,*i*_ as the number of times that a cell was the daughter cell or the parent cell, respectively, in the last *i* elements in the sequence. Accordingly, we define the rate parameter for an arbitrary sequence

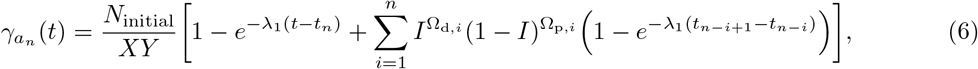

where *t_i_* for 1 ≤ *i* ≤ *n* is obtained from a Beta distribution as defined in (5) and *t*_0_ = 0. We note that there are 2^*n*^ permutations of the cell division sequence, and hence there are 2^*n*^ rate parameters for the cell subpopulation that has experienced *n* divisions and the corresponding mixture distribution has 2^*n*^ components.

We now highlight the validity of this approximate dosage distribution for each cell subpopulation with respect to dosage distribution obtained from the simulation. In Figures 5(a)-(f), we present the dosage distribution for the entire cell population, obtained from the simulation. We observe the asymmetry in the distribution that arises from the asymmetric inheritance process. We next separate the dosage distribution into its constituent components of the dosage distribution for cell subpopulations (Figure 5(g)) that contain cells that have experienced division a specific number of times, and present these distributions for zero, one, and two divisions in Figures 5(h)-(y). As expected, we observe that cells that have experienced fewer divisions are associated to more nanoparticles. Interestingly, while the increase in the mean number of associated nanoparticles per cell is nonlinear, the increase in the mean number of associated nanoparticles per cell for each subpopulation is approximately linear. This implies that the apparent plateau in nanoparticle association can be attributed to cell division rather than dosage depletion effects. We also present the dosage distributions obtained from sampling from the mixture distributions defined via (5)–(6). We observe that the mixture distributions match the simulation results well. However, the approximation of the mixture distribution becomes less accurate as both time and the number of divisions increase. This is because the assumption of exponential growth becomes less appropriate as the number of cells in the population increases. In these results, the average cell doubling time is 12 h. Due to the assumption of exponential waiting times between cell division events, it is possible for a single cell in the model to divide multiple times over a short timeframe. While biologically implausible, the assumption of exponential waiting times between events underpins ordinary differential equation models [46]. As such, the cell division behaviour in our model is consistent with previous models of nanoparticle-cell interactions with cell division [27, 53]. A detailed discussion about the waiting time distributions arising from the connection between stochastic models and ordinary differential equation models can be found in the work by Hurtado and Kirosingh [46]. In Appendix C, we present the fraction of the cell population that have undergone a specific number of divisions by time *t* in the simulation framework under different Π values, that is, for different numbers of stages in the proliferation process. We observe that, as expected, the single stage proliferation model yields results that may not be biologically plausible; however, this is only for a small fraction of the cells in the population. The question of how to model the distribution of times between cell division events is explored in significant depth in the work by Yates *et al*. [42]. The choice of an exponential waiting time facilitates analytic progress but produces a slight overestimation in expected heterogeneity, as noted from the results in Figure 4.

**Figure 5:**
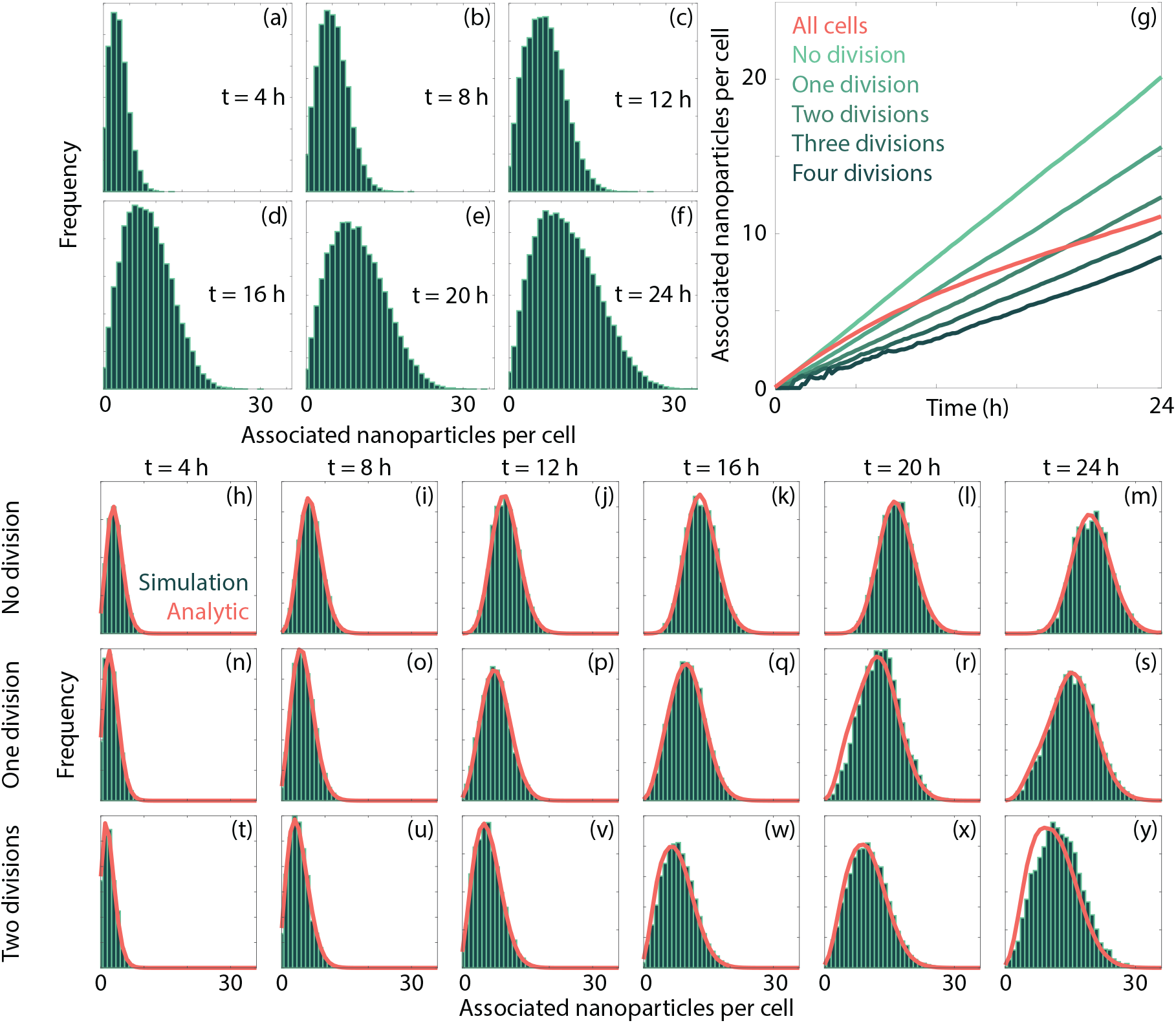
Dosage distributions for subpopulations of cells based on the number of divisions that a cell has experienced. (a)-(f) Dosage distributions recorded at four hour intervals for the entire cell population. (g) Mean number of associated nanoparticles per cell for each cell subpopulation. (h)-(y) Comparison between the analytic (solid line) and simulation (histogram) predictions of the dosage distribution for cells that have experienced (h)-(m) zero, (n)-(s) one, and (t)-(y) two cell divisions recorded at four hour intervals. Parameters used are *d* = 50 nm, *N*_initial_ = 10^5^, *h* = 5 × 10^−3^ m, *λ*_1_ = 8.42 × 10^−4^ h^−1^, 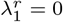, *S* = 1, *C*_0_ = 10, *I* = 0.7, Π = 1, *P*_1_ = 1/72, *K*_1_ → ∞. Analytic predictions are constructed from 10^6^ samples of the mixture distribution. All simulation data are obtained from 1000 identically-prepared realisations of the simulation.

Thus far, we have considered a simplified scenario where the rate of nanoparticle association does not depend on the number of previously-associated nanoparticles. While this is appropriate in certain circumstances, such as for highly phagocytic cell species [24], previous investigations have demonstrated that the nanoparticle association process can become saturated for a sufficiently high administered nanoparticle dose. This could occur for a variety of reasons, including a finite receptor capacity, a finite subcellular compartment capacity, volume constraints within the cell itself, or a balance between nanoparticle association and export. The presence of a carrying capacity would imply that the rate of nanoparticle association decreases with the number of associated nanoparticles [24]. Here saturation is attributed to a finite carrying capacity of individual cells, where there is a maximum number of associated nanoparticles per cell. We note that we investigate other mechanisms that can give rise to nanoparticle saturation elsewhere in this manuscript. To capture the effect of a cell carrying capacity, we now apply a finite *K*_1_ value, which enforces a decrease in the effective rate parameter 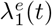 as the number of associated nanoparticles increases. We present the dosage distributions obtained from the simulation with a finite carrying capacity in Figure 6. We observe that at early times, where the mean number of associated nanoparticles is small compared to the carrying capacity, the dosage distribution is approximately Poisson distributed (Figure 6(a)). However, as the mean number of associated nanoparticles approaches the carrying capacity, we observe that the dosage distribution becomes primarily located near the carrying capacity (Figure 6(f)). As such, the assumption of a Poisson dosage distribution is not always appropriate, as observed previously [8]. In Figure 6(g) we see that the standard deviation of the dosage distribution decreases as the mean number of associated nanoparticles approaches the carrying capacity, as opposed to the monotone increase that is expected if the dosage is Poisson distributed. This highlights the benefit of employing the modelling framework to generate predictions of the dosage distribution, as it is not always possible to rely on analytic results, and the inclusion of specific biological mechanisms can influence the features of the dosage distribution.

**Figure 6:**
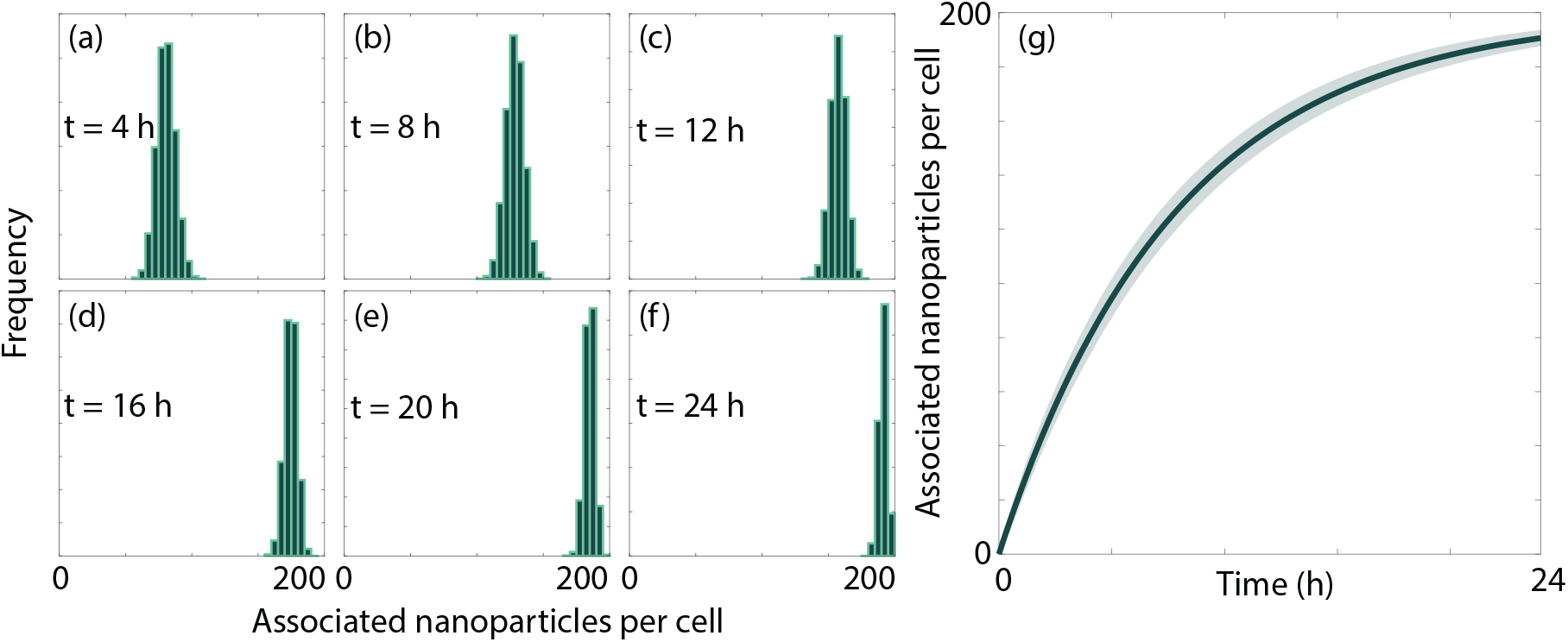
Dosage distributions for cells with a finite carrying capacity of nanoparticles. (a)-(f) Dosage distributions recorded at four hour intervals obtained from the simulation. (g) Mean (± one standard deviation) number of associated nanoparticles per cell obtained from the simulation. Parameters used are *d* = 50 nm, *N*_initial_ = 2 × 10^6^, *h* = 3 × 10^−3^ m, *λ*_1_ = 1.27 × 10^−3^ h^−1^, 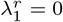, *S* = 1, *C*_0_ = 100, *K*_1_ = 200. All simulation data are obtained from 1000 identically-prepared realisations of the simulation.

We now consider the impact of nanoparticle recycling on the dosage distribution, which may also induce saturation in the number of associated nanoparticles. We implement this in our model, by allowing associated nanoparticles to return to the media with rate 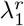. We present the dosage distributions and mean number of associated nanoparticles per cell in Figure 7. We observe that a plateau in the number of associated nanoparticles per cell occurs, similar to that observed in the case with either dosage depletion or a carrying capacity. The plateau arises when the rate of nanoparticle association is balanced by the rate of nanoparticle recycling. However, unlike the carrying capacity case, the dosage distribution remains Poisson distributed. This can seen by comparing the standard deviation from the simulations against the standard deviation from the corresponding Poisson distribution, as highlighted in Figure 7(g).

**Figure 7:**
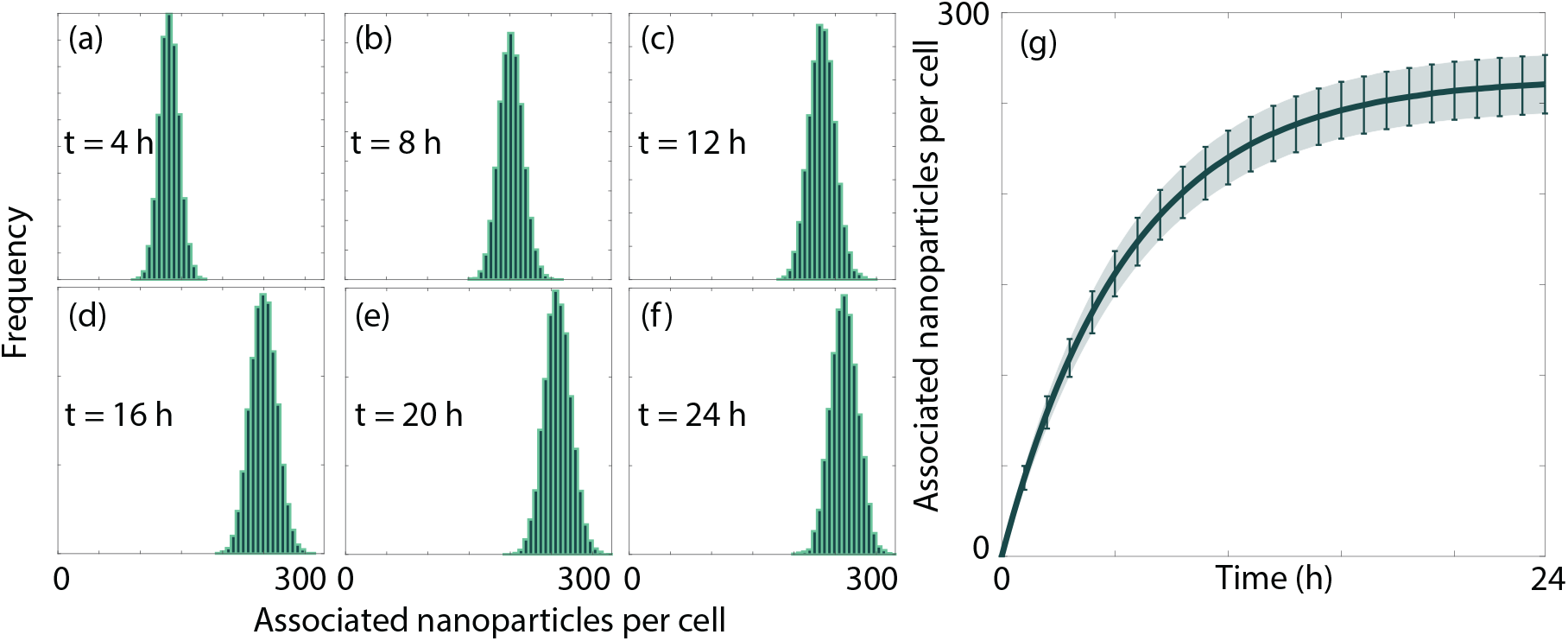
Dosage distributions for cells with nanoparticle recycling. (a)-(f) Dosage distributions recorded at four hour intervals obtained from the simulation. (g) Mean (± one standard deviation) number of associated nanoparticles per cell obtained from the simulation. The ribbon is the standard deviation obtained from the simulation, whereas the error bars are the standard deviation predicted by a Poisson distribution with the mean value obtained from the simulation. Parameters used are *d* = 50 nm, *N*_initial_ = 5 × 10^5^, *h* = 5 × 10^−4^ m, *λ*_1_ = 9.30 × 10^−3^ h^−1^, 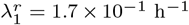, *S* = 1, *C*_0_ = 100, *K*_1_ → ∞. All simulation data are obtained from 1000 identically-prepared realisations of the simulation.

### 3.2. Nanoparticle binding and internalisation

To now, we have focused on a single stage model of nanoparticle internalisation. In reality, the nanoparticle internalisation process is significantly more complicated than can be captured by a single stage model [23]. The reason such simple models see widespread usage is that it is not standard practice to measure the relevant experimental information necessary to distinguish between the stages of internalisation, and hence more complicated models cannot always be parameterised. Further details can found in the review of Fitzgerald and Johnston [19], who discuss the benefits and drawbacks of the various experimental techniques that can be used to distinguish between nanoparticles that are strongly bound to a cell’s membrane, and those that have been internalised. However, it is still instructive to consider how the number of nanoparticles in each stage of the internalisation process changes over time, under certain assumptions.

We next consider an irreversible two stage model, where nanoparticles first bind to a cell’s membrane and, subsequently, become internalised by that cell. We assume that the cell population is fully confluent and that the carrying capacity at each stage of internalisation can be neglected. For a nanoparticle to be bound to a cell at time *t*, we require that the time of binding is less than *t* and that the time of internalisation is greater than *t*. If these two events occur according to two exponential random variables *X*_1_ and *X*_2_ with rates *λ*_1_ and *λ*_2_, then the probability that an individual nanoparticle is bound at time *t* is equivalent to *P* (*X*_1_ < *t, X*_1_ + *X*_2_ > *t*). Accordingly, the mean number of bound nanoparticles per cell is

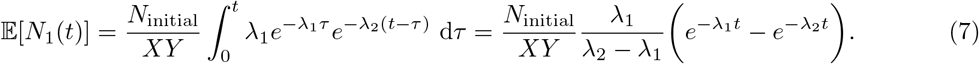

For a nanoparticle to be internalised at time *t*, we require that the combined binding and internalisation time is less than *t*, that is, *P* (*X*_1_ + *X*_2_ < *t*). Hence the mean number of internalised nanoparticles per cell is

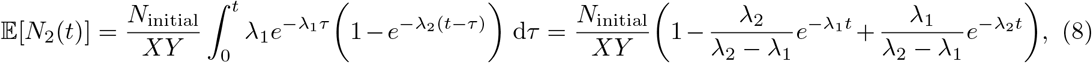

which is the CDF of a hypoexponential random variable with parameters *λ*_1_ and *λ*_2_ [54], multiplied by the average number of nanoparticles initially present in the culture media above each lattice site. The dosage distributions for bound and internalised nanoparticles, under the aforementioned assumptions, are Poisson distributions with rate parameters 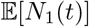 and 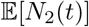, respectively. These results can be extended to describe cell populations that are not at confluence, where

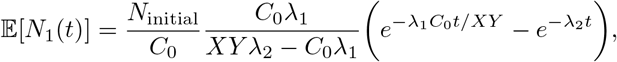

and

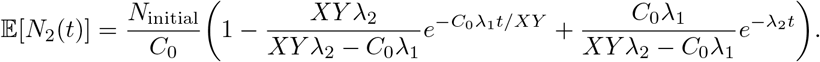

As for the single stage internalisation model, the effective rate parameter for binding decreases with confluence, but this is countered by the increase in the number of nanoparticles available to an individual cell, as the total number of nanoparticles in the culture media is constant.

We present a comparison between the analytic and simulation predictions of the dosage distribution for associated, bound, and internalised nanoparticles in Figure 8. The number of associated nanoparticles is the sum of the number of bound and internalised nanoparticles. We observe that the analytic predictions accurately describe the dosage distribution arising from the simulation, both in the case where the cell population is fully confluent (Figure 8(a)) and where the cell population is not fully confluent (Figure 8(b)). Further, these results highlight the importance of accurately distinguishing between bound and internalised nanoparticles [37], rather than reporting the number of associated nanoparticles. We see similar trends in nanoparticle association between Figures 8(a) and 8(b). However, the ratio of bound to internalised nanoparticles varies substantially due to the difference in the ratio of the binding and internalisation rate parameters. If the effective treatment of a cell relies on a threshold dose of nanoparticles being internalised, then we may observe a distinct response to the treatment, even though the number of associated nanoparticles is similar.

**Figure 8:**
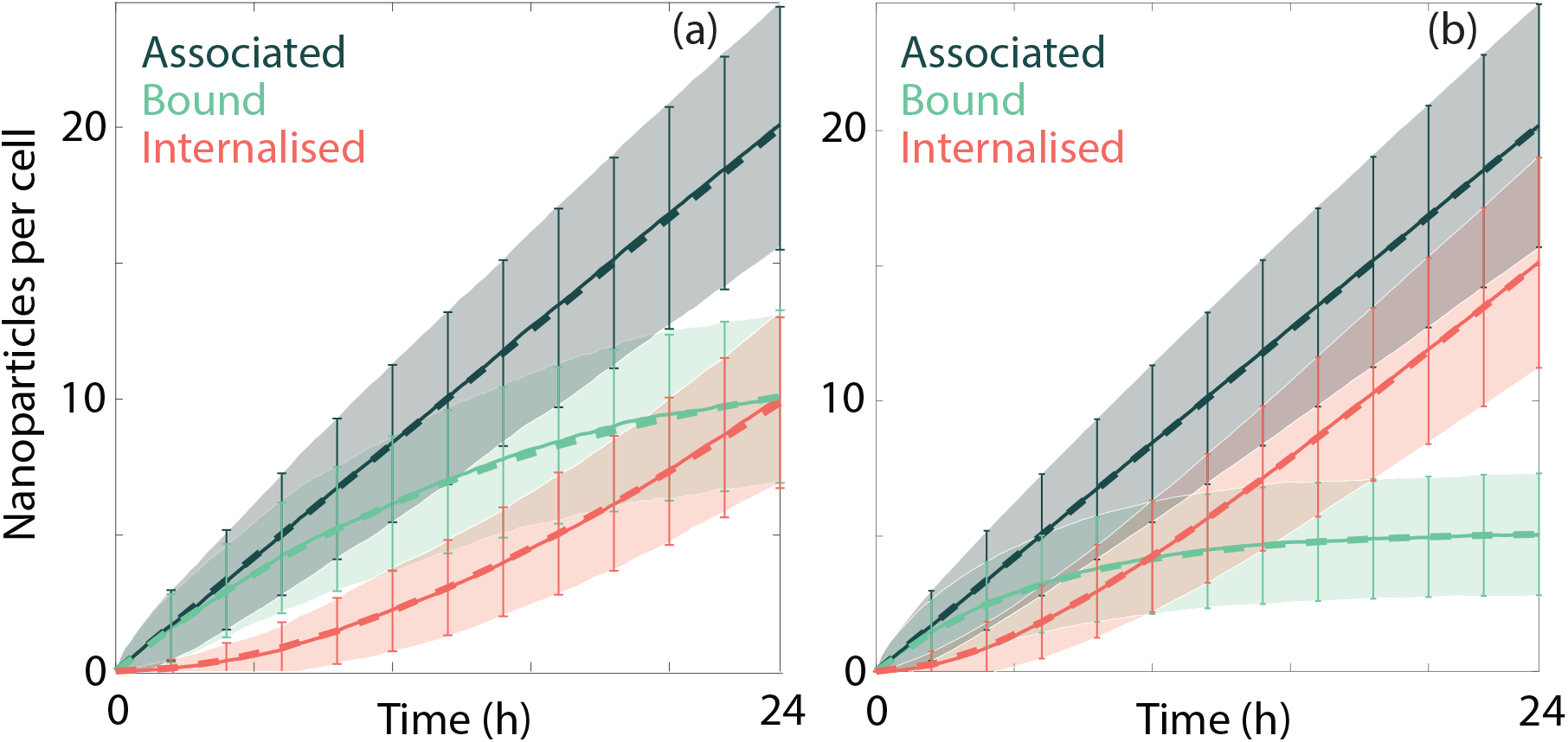
Comparison between the analytic and simulation predictions of the number of associated (dark green), bound (light green) and internalised (red) nanoparticles per cell at (a) full confluence and (b) 10% confluence. Lines correspond to the analytic (dashed) and simulation (solid) predictions of the mean. The analytic and simulation predictions of the mean ± one standard deviation correspond to the error bars and ribbons, respectively. Parameters used are *S* = 2, *K*_1_ → ∞, *K*_2_ → ∞, *N*_initial_ = 10^5^, *λ*_1_ = 8.40 × 10^−4^ h^−1^, 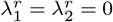, (a): *d* = 50 nm, *h* = 5 × 10^−3^ m, *λ*_2_ = 6.49 × 10^−2^ h^−1^, *C*_0_ = 100; (b): *P_p_* = 0 ∀ *p*, *d* = 20 nm, *h* = 3×10^−3^ m, *λ*_2_ = 1.62 × 10^−1^ h^−1^, *C*_0_ = 10. All simulation data are obtained from 500 identically-prepared realisations of the simulation.

It is possible to calculate the mean number of nanoparticles per cell for each stage of the internalisation process for an arbitrary number of stages in the absence of any carrying capacities. While we state the result here, we note that it may not be possible to parameterise a model with more than two stages from the types of experimental data that are routinely collected, and hence we do not focus on analysing simulation results for *S* > 2 here. The number of nanoparticles that have reached the *S*th stage of internalisation (i.e. that have been internalised) at time *t* is

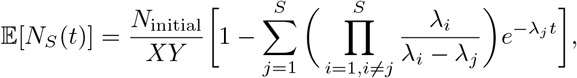

which arises from an *S*-stage hypoexponential distribution [54]. We can use this result to recursively define the number of nanoparticles in the *s*th stage of internalisation at time *t* for 1 ≤ *s* ≤ *S* − 1, as the number of nanoparticles in the *s*th stage is the difference between the number of nanoparticles that have reached the *s*th and (*s* + 1)th stages of internalisation

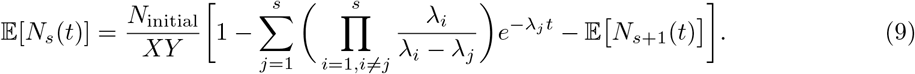

We note that we can describe cell populations that are not fully confluent via a rescaling of *λ*_1_ and the number of nanoparticles available to each cell by *XY/C*_0_ as previously. In the absence of a carrying capacity and cell proliferation, the dosage distribution for the *s*th stage of internalisation is a Poisson distribution with rate parameter 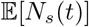.

If cell proliferation cannot be neglected, a Poisson dosage distribution is no longer appropriate. As in the case of the single stage internalisation model, it is possible to predict the dosage distribution by considering subpopulations of cells based on the number of division events that a cell has experienced. Again, we require that dosage depletion effects can be neglected, and that the initial confluency of the cell population is small, such that cell division can approximated via exponential growth. The distribution of cell division times is Beta distributed as defined in (5). For cells that have divided exactly once, the distribution of the number of bound nanoparticles per cell is a mixture distribution of equally-weighted two Poisson distributions with rate parameters

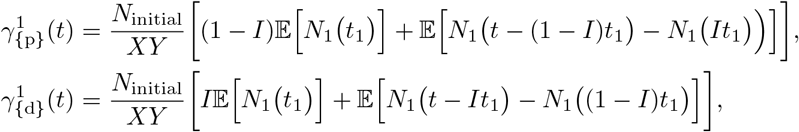

where *t*_1_ ~ *U* (0, *t*), 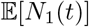 is defined in Eqauation (7), and the superscript denotes that the rate parameters correspond to the first stage of the internalisation process. The distribution of the number of internalised nanoparticles per cell is a mixture distribution of two equally-weighted Poisson distributions with rate parameters

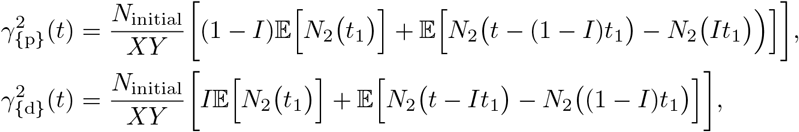

where *t*_1_ ~ *U* (0, *t*), 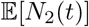 is defined in Eqauation (8), and the superscript denotes that the rate parameters correspond to the second stage of the internalisation process.

We can express the components of the mixture distribution that represents the dosage of nanoparticles in the *s*th stage of internalisation for the subpopulation of cells that have experienced *n* divisions in general terms. Recall that *a_n_* = {*b*_1_, *b*_2_, · · ·, *b_n_*}, where *b_j_* ∈ {d,p}, defines the number, order, and type of division events that a cell has experienced. The mixture distribution for the subpopulation of cells that have experienced exactly *n* divisions has 2^*n*^ equally-weighted components, each of which is a Poisson distribution with rate parameter

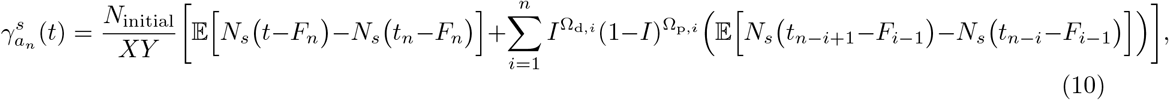

where

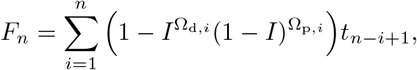

is the time shift that is proportional to the amount of nanoparticles [lost to]division, *t_i_* for 1 ≤ *i* ≤ *n* is obtained from a Beta distribution as defined in (5), *F*_0_ = 0, *t*_0_ = 0, 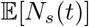 is defined in Eqauation (9), Ω_d,*i*_ and Ω_p,*i*_ are the number of times that a cell was the daughter cell or the parent cell, respectively, in the last *i* elements in the sequence *a_n_*. We include the *F_n_* terms here as the number of nanoparticles in each stage can decrease due to nanoparticles progressing through the internalisation pathway for the multistage internalisation model. We could include these terms in the single stage model; however, as there are no additional stages of internalisation, and we are in the linear association regime, it will have no effect. While this mixture distribution is not presented in a closed form, it allows us to efficiently sample from the dosage distribution at a particular time without performing realisations of the simulation.

We present a comparison of the mixture distribution and the dosage distribution arising from the simulation for a two stage internalisation model in Figure 9. We first present the mean number of bound and internalised nanoparticles per cell in Figures 9(a)-(b), respectively, both for the entire cell population, and for cell subpopulations separated based on the number of division events experienced. Again, we observe that cells that have experienced more division events have fewer bound and internalised nanoparticles. We present the dosage distributions for bound and internalised nanoparticles for cells that have experienced zero, one and two divisions in Figures 9(i)-(z). We see that the mixture distributions accurately approximate the dosage distributions obtained from the simulation. There are minor discrepancies between the mixture distributions and the simulation dosage distributions that appear as the number of cells in the population increases and the approximation of exponential growth is less accurate. However, as obtaining a sample from the mixture distribution for cells that have experienced *n* divisions only requires sampling *n* + 1 random variables, it is significantly more efficient to use the mixture distributions to predict the dosage distribution across each of the cell subpopulations.

**Figure 9:**
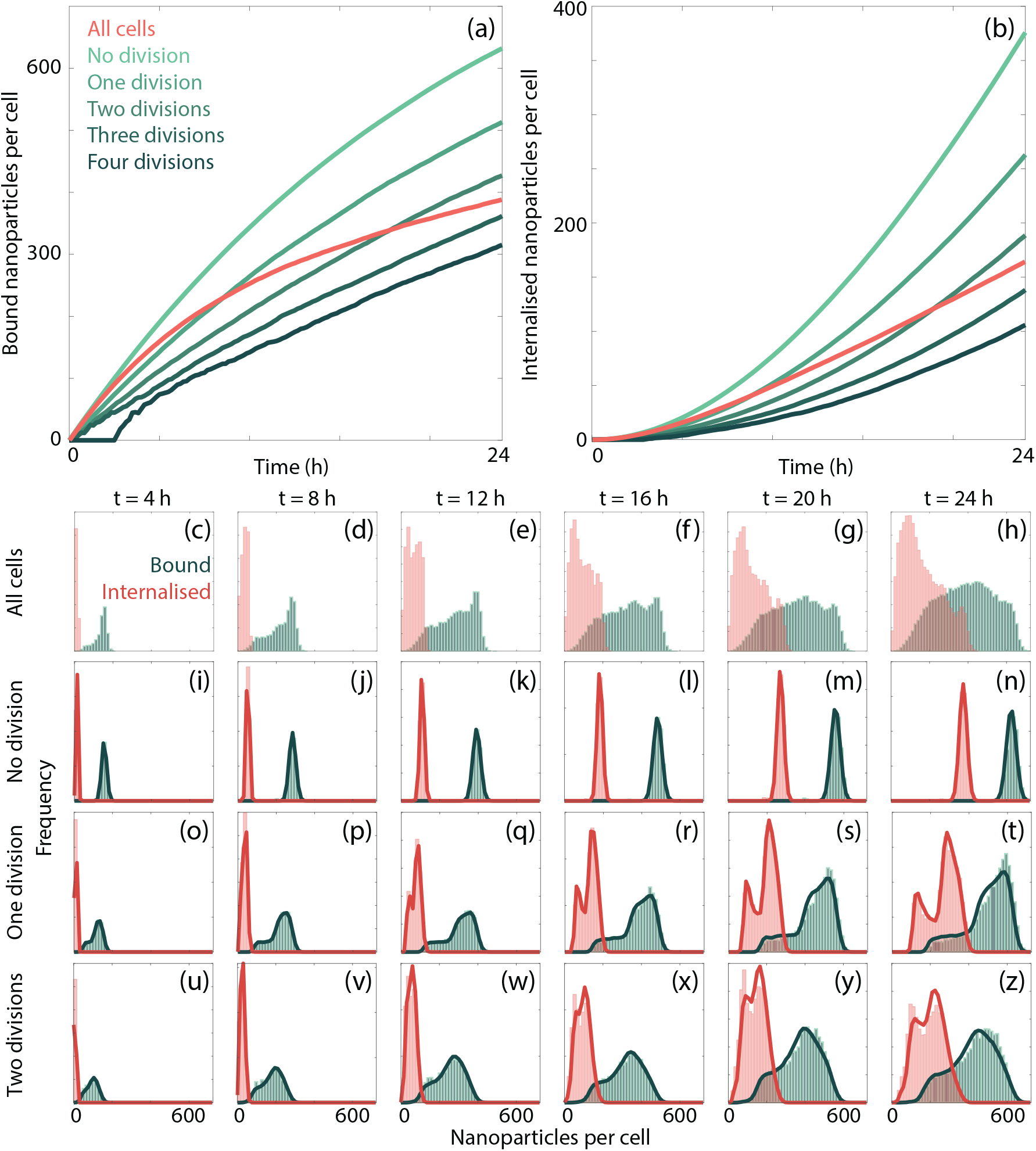
Dosage distributions of bound and internalised nanoparticles for subpopulations of cells based on the number of divisions that a cell has experienced. (a)-(b) Mean number of (a) bound and (b) internalised nanoparticles per cell for each cell subpopulation. (c)-(h) Dosage distributions for bound (green) and internalised (red) nanoparticles recorded at four hour intervals for the entire cell population. (i)-(z) Comparison between the analytic (solid line) and simulation (histogram) predictions of the dosage distribution for bound (green) and internalised (red) nanoparticles for cells that have experienced (i)-(n) zero, (o)-(t) one, and (u)-(z) two cell divisions recorded at four hour intervals. Parameters used are *d* = 50 nm, *N*_initial_ = 5 × 10^6^, *h* = 5 × 10^−3^ m, *S* = 2, *λ*_1_ = 8.42 × 10^−4^ h^−1^, *λ*_2_ = 4.26 × 10^−2^, 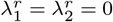, *C*_0_ = 10, *I* = 0.7, Π = 1, *P*_1_ = 1/72, *K*_1_ → ∞. Analytic predictions are constructed from 10^6^ samples of the mixture distribution. All simulation data are obtained from 1000 identically-prepared realisations of the simulation.

Previous investigations have highlighted the apparent saturation of nanoparticle association [23, 24, 27, 28]. However, if this can be attributed to a finite cell carrying capacity, it is not obvious which stages of the internalisation process cause this saturation. For example, it is possible that a cell expresses a finite number of receptors that nanoparticles bind to in sufficient numbers such that there are no free receptors for further nanoparticle binding. Alternatively, cells may have a finite internal volume available to be filled by nanoparticles, after which no further internalisation may occur, which may explain the saturation in nanoparticle association. To account for these possibilities we now introduce a finite carrying capacity for each stage of the internalisation process.

We consider two possible scenarios to examine the influence of finite carrying capacities. First, we consider the case where the rates of both nanoparticle binding and nanoparticle internalisation decrease with the number of previously bound and internalised nanoparticles, respectively. We present the dosage distributions for bound and internalised nanoparticles, obtained from the simulation, as well as the mean number of associated, bound and internalised nanoparticles per cell in Figures 10(a)-(g). Here we select *K*_1_ = 100 and *K*_2_ = 50, that is, at most 100 nanoparticles can be bound to an individual cell and, at most, 50 nanoparticles can be internalised by a cell. We see that the carrying capacities result in dosage distributions that are not Poisson, particularly as the number of bound/internalised nanoparticles approaches the relevant carrying capacity. As expected, the mean number of bound and internalised nanoparticles per cell both plateau as the carrying capacities are approached, which is reflected in a corresponding plateau in the mean number of associated nanoparticles per cell. In the second case, only the rate of nanoparticle binding decreases with the number of previously bound nanoparticles. Here we choose *K*_1_ = 100 and *K*_2_ → ∞. In the dosage distributions obtained from the simulation, presented in Figures 10(h)-(n), we see that the level of asymmetry in the number of bound nanoparticles is reduced, compared to the case with two carrying capacities. This is because it is always possible for a nanoparticle to transition from bound to internalised, and hence we do not see a buildup in the bound nanoparticle dosage distribution located at the carrying capacity. The dosage distribution for the internalised nanoparticles is approximately symmetric, and is more consistent with dosage distributions obtained in the absence of any carrying capacities. Interestingly, while we see a distinct plateau in the number of bound nanoparticles per cell, the fact that the number of internalised nanoparticles per cell is free to increase means that the number of associated nanoparticles per cell continues to increase. There is a qualitative difference in the shape of the nanoparticle association curves: the presence of both carrying capacities results in a plateau, compared to a single carrying capacity for the bound nanoparticles, which results in an initial decrease in the rate of association as the number of bound nanoparticles per cell plateaus, followed by a constant increase in the number of associated nanoparticles per cell. This implies that it may be possible to develop an experimental protocol that distinguishes between scenarios where carrying capacities may or may not be relevant at different stages of internalisation, even if the number of bound and internalised nanoparticles are not distinguished in the experimental data.

**Figure 10:**
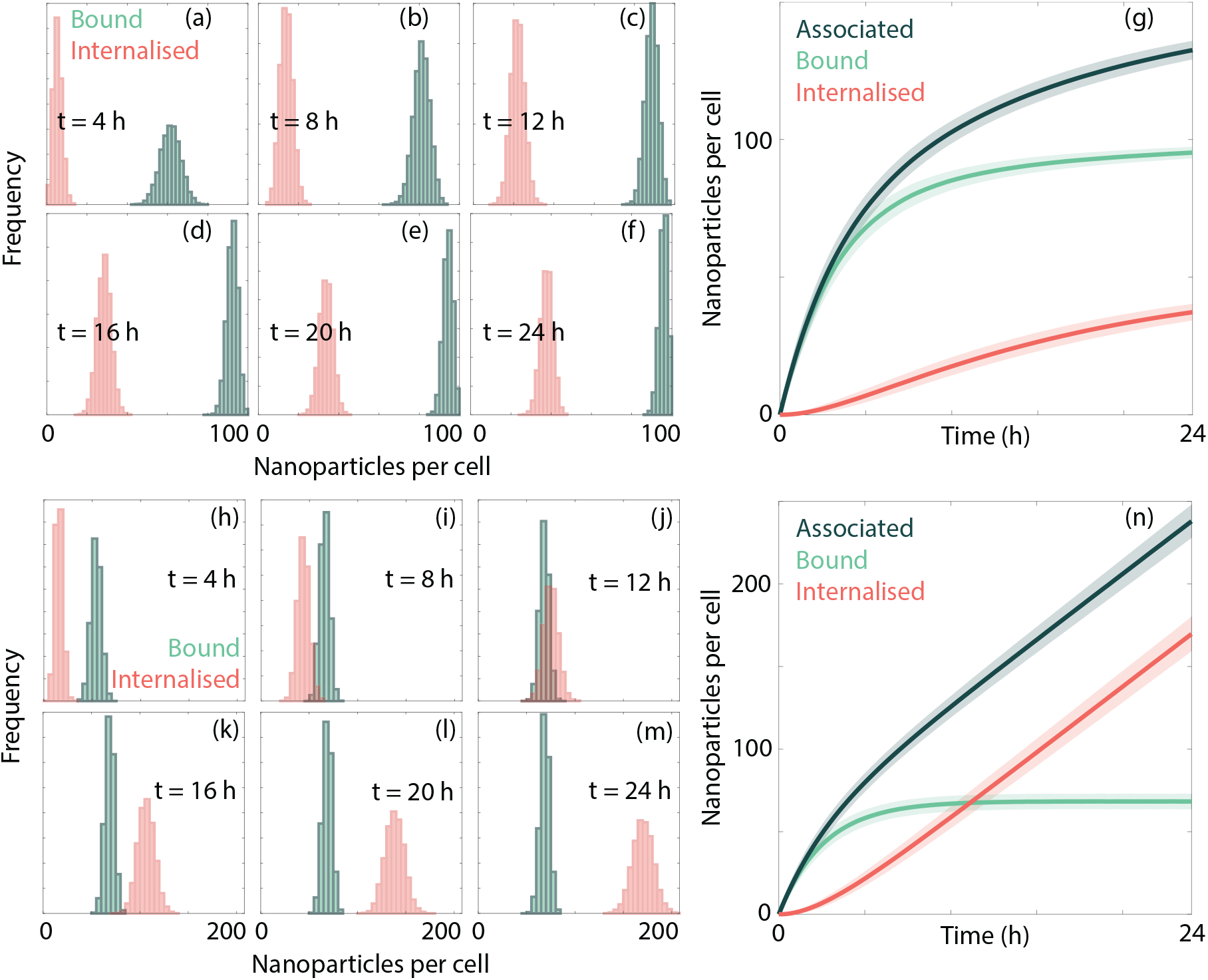
Comparison of nanoparticle internalisation with (a)-(g) a finite carrying capacity for both bound and internalised nanoparticles and (h)-(n) a finite carrying capacity for bound nanoparticles only. (a)-(f), (h)-(m) Dosage distributions recorded at four hour intervals for bound (green) and internalised (red) nanoparticles for cells with a finite carrying capacity of nanoparticles, obtained from the simulation. (g),(n) Mean (± one standard deviation) number of associated (dark green), bound (light green) and internalised (red) nanoparticles per cell obtained from the simulation. Parameters used are *d* = 50 nm, *N*_initial_ = 2 × 10^6^, *h* = 3 × 10^−3^ m, *S* = 2, *C*_0_ = 100,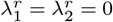, (a)-(g): *λ*_1_ = 1.27 × 10^−3^ h^−1^, *λ*_2_ = 3.62 × 10^−2^ h^−1^, *K*_1_ = 100, *K*_2_ = 50; (h)-(n): *λ*_1_ = 1.27 × 10^−3^ h^−1^, *λ*_2_ = 1.17 × 10^−1^ h^−1^, *K*_1_ = 100, *K*_2_ → ∞. All simulation data are obtained from 1000 identically-prepared realisations of the simulation.

### 3.3. Model identification and parameter estimation

The modelling framework presented here can provide predictions about heterogeneity in nanoparticle dosage, based on assumptions about the biological mechanisms that are relevant for a particular experimental design. However, the questions of whether the presence of key biological mechanisms can be identified from standard experimental data [19], and the corresponding parameters estimated, remain open. While these questions merit an in-depth investigation, here we highlight certain cases where parameter estimation is possible, via both heuristic and numerical approaches. These approaches are informed by the form of the analytic results presented here; though we stress parameter estimation is not the primary aim of this manuscript. Further, we identify cases where it is not possible to distinguish between putative biological mechanisms using standard experimental data, motivating potential changes to experimental protocols.

A key parameter that informs the efficacy of a potential nanoparticle-based therapeutic is the nanoparticle-cell affinity [24]. This parameter provides a quantitative estimate of the strength of interaction between nanoparticles and cells [24]. In the context of the model presented here, the nanoparticle-cell affinity is proportional to *λ*_1_. While this parameter can be estimated for each case where an analytic result is presented via numerical nonlinear curve fitting methods, provided that 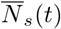 is measured for all *s*, where the bar indicates that this is the mean of the experimental data, such methods can require specialist knowledge to implement. Although such an approach would be preferable, here we discuss heuristic approaches for estimating key biological parameters, which require less specialised knowledge. For example, from the result in Equation (1), we know that at early experimental time points we should observe that the rate of nanoparticle association is approximately linear, with the caveat that there is negligible cell division and a high carrying capacity. Accordingly, *λ*_1_ can simply be estimated via the slope of nanoparticle association over time. If the experiment is performed for sufficiently long, and there is reason to include a cell carrying capacity, *K*_1_ can be estimated via inspection. The estimation approach can be extended for multistage internalisation processes, as *λ*_1_ can still be estimated from nanoparticle association data. Provided that 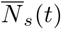 is measured for all *s* and we have an estimate of *λ*_1_, other transition rates can be estimated heuristically from the data. For example, for a two-stage internalisation process, we can estimate

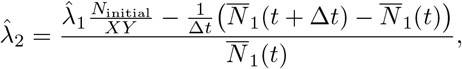

where 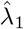 and 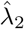 are the estimated internalisation rate parameters. The estimate 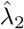 is obtained from the expected change in the number of bound nanoparticles between observations captured at a time Δ*t* apart. Similar processes can be followed to estimate the rate parameters for other internalisation stages, provided the appropriate 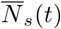 is measured experimentally. However, we note that this approach relies on derivatives estimated from experimental data via finite difference approximations, which may not be appropriate in the presence of substantial experimental noise [55].

We now demonstrate that the simulation model can be compared against experimental data with parameters that are estimated heuristically. However, we add the caveat that here we rely on relatively strong assumptions about the presence of specific mechanisms to be able to make this comparison. We consider previously-published data for 150 nm and 214 nm (poly)methacrylic acid nanoparticles and RAW 264.7 macrophages [24]. Additional details are included in Appendix A. For the results in Figures 11(a)-(g), guided by the previous experimental investigation into the data [24], we assume that there is a finite carrying capacity, that cell division is not significant, and that nanoparticle recycling does not play a role. For the results in Figures 11(h)-(m), again guided by the previous investigation [24], we assume that cell division is not significant, that nanoparticle recycling does not play a role, and that there is no finite carrying capacity. It is obviously preferable to not rely on such strong assumptions, and hence the results here should only be treated as a proof of concept that highlight the presence of biological heterogeneity consistent with previous investigations [8, 9]. An in-depth investigation into the types of experimental data necessary to robustly determine the presence and strength of each of the potential mechanisms considered in this manuscript would be instructive. However, such an investigation is well beyond the scope of this manuscript.

**Figure 11:**
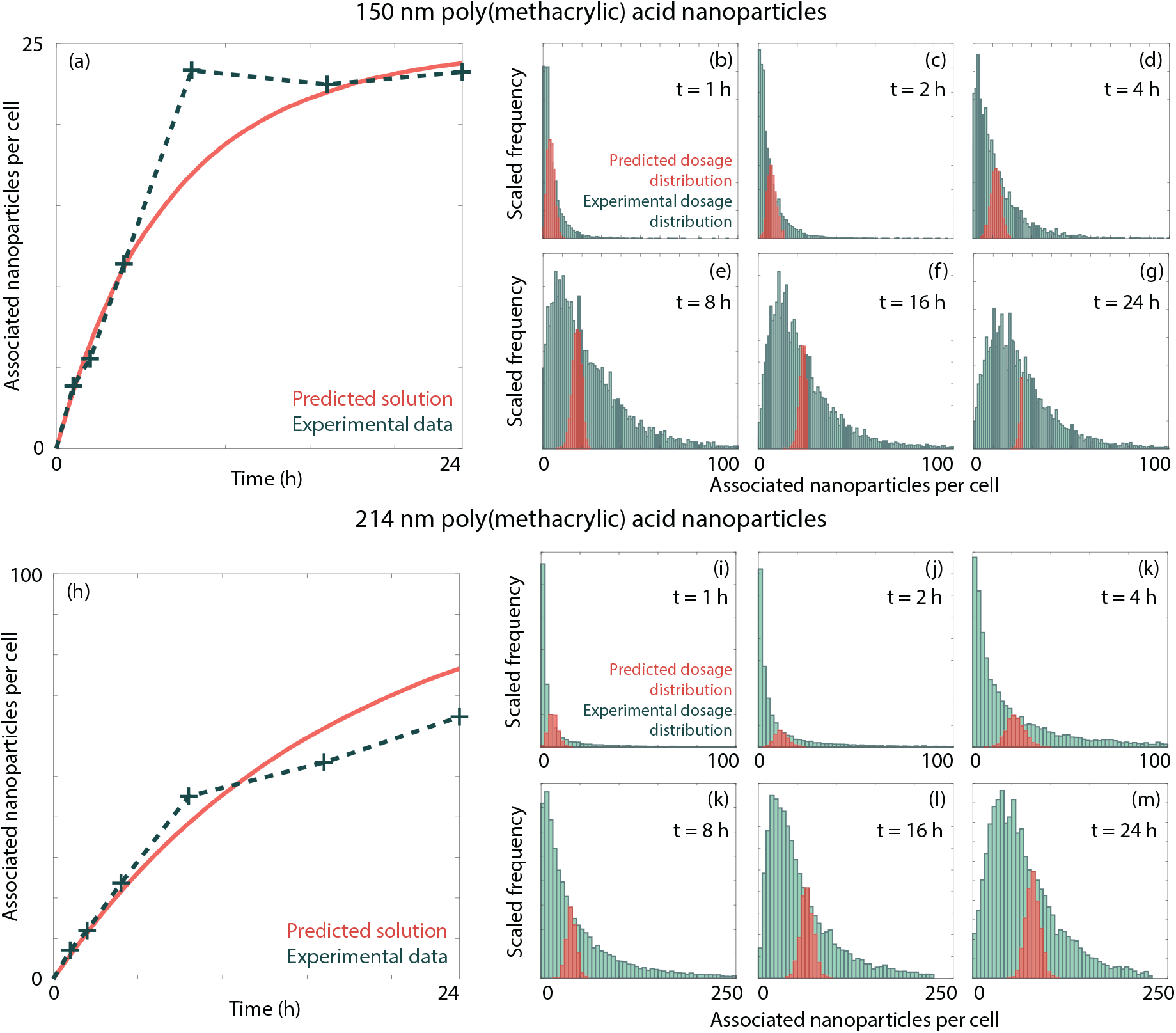
Comparison between the simulation model and experimental data via heuristic parameter estimation. (a)-(g) Comparison between the simulation data model (red) and experimental data (green) for the (a) average number of associated nanoparticles per cell and (b)-(g) dosage distributions for 150 nm (poly)methacrylic acid nanoparticles and RAW 264.7 macrophages. (h)-(m) Comparison between the (h) average number of associated nanoparticles per cell and (i)-(m) dosage distributions for 214 nm (poly)methacrylic acid nanoparticles and RAW 264.7 macrophages. Parameters used are *N*_initial_ = 10^5^, *h* = 0.5 × 10^−3^ m, *S* = 1, *C*_0_ = 100, 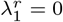, (a)-(g): *λ*_1_ = 4.78 × 10^−2^ h^−1^, *K*_1_ = 25, *d* = 150 nm; (h)-(m) *λ*_1_ = 5.99 × 10^−2^ h^−1^, *K*_1_ → ∞, *d* = 214 nm. The frequencies are scaled such that the mode of the predicted dosage distribution is equal to the value of the experimental dosage distribution at that dosage.

Under the above assumptions, we can estimate the relevant parameters in the model from the data, and then compare the predicted dosage distributions against the experimental dosage distributions. We use the observations at *t* = 0, 1, 2 h to estimate *λ*_1_. We present a comparison between the average number of associated nanoparticles obtained from the experiment and from the simulation in Figures 11(a),(h). In both cases, we see that the results from the simulation match the experimental data. We next turn to the comparison of the dosage distributions, as presented in Figures 11(b)-(g) and 11(i)-(m). We observe that the experimental dosage distributions are significantly wider than can be attributed to the stochastic nature of nanoparticle association, a result consistent with previous investigations [8, 9]. This suggests that the underlying cell population exhibits considerable biological heterogeneity. In particular, the assumption of a finite carrying capacity reduces the width of the dosage distribution obtained from the simulation, which is not observed in the experimental. This may imply the presence of a carrying capacity that varies between cells [8] or that the plateau in nanoparticle association is attributable to other factors. However, as noted above, here we require relatively strong assumptions to compare the model with the experimental data. This highlights the need for an indepth investigation to determine experimental protocols that allow for these assumptions to be relaxed.

To reinforce this point, we highlight pitfalls that can occur when attempting to identify the biological mechanisms that dictate nanoparticle internalisation from nanoparticle association assay data. We present the mean number of associated nanoparticles per cell for three scenarios in Figure 12. The first scenario is where the cell population has a finite carrying capacity. The second scenario is where there is no carrying capacity, but the nanoparticles associate sufficiently rapidly over the course of the experiment that dosage depletion effects are relevant. The third scenario is where there is no carrying capacity or dosage depletion, but nanoparticle recycling occurs. As we observe in Figure 12, it is difficult to distinguish between the mean number of associated nanoparticles per cell for these three scenarios, as parameters can be selected such that each case exhibits a similar plateau. It is therefore important to ensure that the experiment protocol allows for the association to be measured relative to the initial dosage, such as by converting fluorescence to nanoparticles per cell [31, 37]. This information would allow for dosage depletion effects to be ruled out; though equally the experiment could be repeated with a higher initial dosage. Similarly, it would be instructive to perform a “pulse and chase” experiment, where nanoparticles only associate to cells at the start of the experiment, and any subsequent nanoparticle recycling can be observed [27]. For example, in the work by Salvati *et al*. [27], the authors attribute a decrease in internalised nanoparticles solely to cell division, rather than any recycling of nanoparticles located in lysosomes. In the absence of these additional measurements, it is not straightforward to say whether a particular mechanism leads to the observed plateau in associated nanoparticles. While we see a difference in the width of the dosage distribution between the case with a finite carrying capacity and the other two cases, it is unclear whether this would be observable when the dosage distribution is confounded by experimental measurement noise and biological heterogeneity. As such, it is critical to follow experimental protocols that allow for quantification of the relevant experimental measurements, and hence allow for reliable estimation of the biological parameters that dictate nanoparticle internalisation [37, 56, 57].

**Figure 12:**
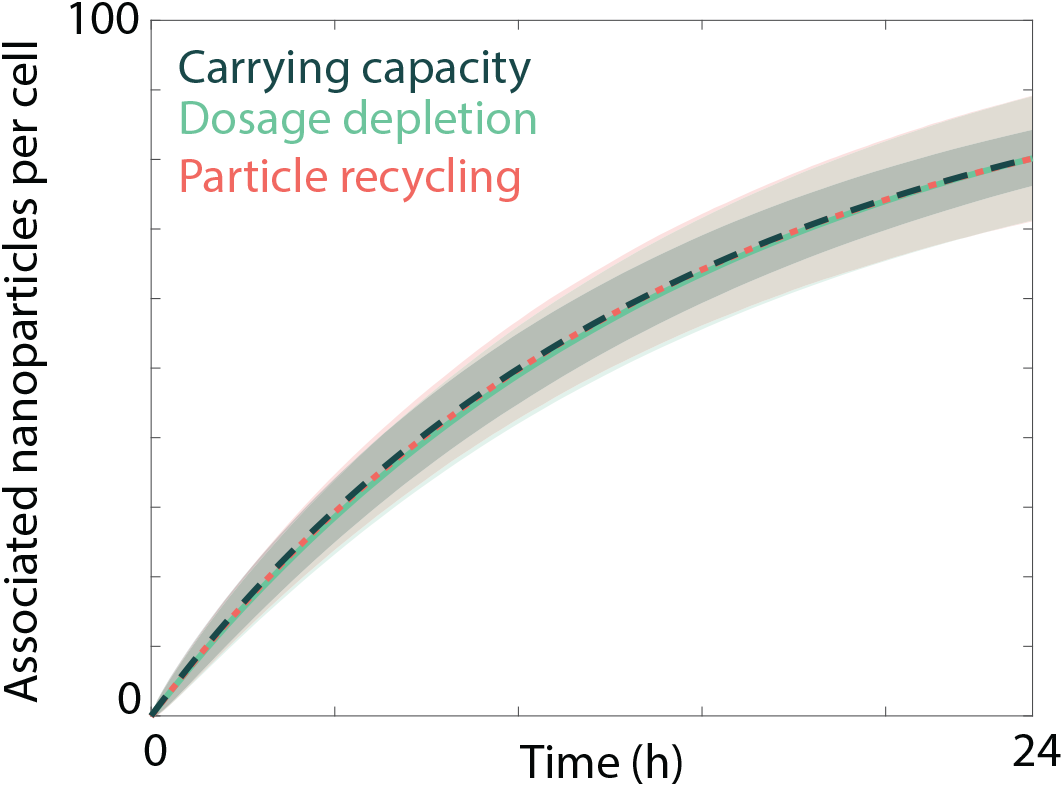
Comparison between the number of associated nanoparticles per cell obtained from the simulation for three potential saturating mechanisms: a finite carrying capacity (dark green, dashed), depletion of nanoparticles in the culture media (light green, solid), and nanoparticle recycling (red, dots). Ribbons correspond to the mean ± one standard deviation. Parameters used are *d* = 20 nm, *h* = 5 × 10^−4^ m, *S* = 1, *C*_0_ = 100, (light green): *λ*_1_ = 6.71 × 10^−2^,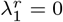, h^−1^, *N*_initial_ = 10^4^, *K*_1_ → ∞; (dark green): *λ*_1_ = 7.075 × 10^−3^ h^−1^, 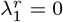, *N*_initial_ = 10^5^, *K*_1_ = 100; (red): *λ*_1_ = 6.94 × 10^−3^ h^−1^, 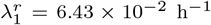, *N*_initial_ = 10^5^, *K*_1_ → ∞. All simulation data are obtained from 200 identically-prepared realisations of the simulation.

## 4. Discussion and conclusions

Nanoparticles are a promising avenue for the targeted delivery of therapeutics to specific cell types [1, 2, 4]. While this technology has seen recent success in specific areas [1], significant gaps remain in the understanding of nanoparticle-cell interactions. This hinders the effective development of nanoparticle-based therapeutics [2, 4, 5, 6]. One factor that complicates analysis and interpretation of standard experiments is heterogeneity in experimental data [8, 9, 10, 11, 12]. It is not straightforward to determine whether this heterogeneity arises from the underlying cell population, or from the stochastic nature of nanoparticle motion and nanoparticle-cell interactions [8, 9]. It is critical to identify whether significant biological heterogeneity is present, as it may imply the existence of a subpopulation of cells that do not respond to a particular treatment [16, 17]. Mathematical models have been employed to investigates the sources of heterogeneity previously [8, 9, 10, 11]; however, such investigations have relied on simplified models of nanoparticle-cell interactions, and neglect the potential impact of a range of biological processes.

Here we present a mathematical modelling framework that provides predictions of the level of heterogeneity expected to manifest in nanoparticle dosage due to the presence of specific biological mechanisms. As such, if we know that particular biological mechanisms are relevant, we are able to identify how much heterogeneity in dosage is unavoidable due to the stochastic nature of the nanoparticle internalisation process. Any additional heterogeneity can therefore be attributed to either experimental measurement error, which is well-established for techniques such as flow cytometry [58], or to heterogeneity in the underlying cell population [8, 9]. We include and investigate a range of biological mechanisms that can be relevant for standard nanoparticle-cell experiments, including multistage nanoparticle internalisation, cell division, asymmetric nanoparticle inheritance and saturation of nanoparticle internalisation. We provide analytic expressions for the expected dosage distributions under certain simplifying assumptions. We demonstrate that in the presence of cell proliferation, the overall dosage distribution can be accurately approximated via a series of mixture distributions for the cell subpopulations that have experienced a specific number of cell divisions. A summary of the key results for each biological mechanism investigated, and whether analytic results can be obtained in each case, is presented in Table 1. The analytic results provide an efficient method for determining the expected nanoparticle dosage distribution. However, as noted throughout, the analytic results rely on simplifying assumptions and hence the conclusions should be interpreted in the context of the model assumptions.

**Table 1:** Summary of the impact of specific biological mechanisms on the nanoparticle dosage distribution, and whether analytic results can be used to calculate the corresponding dosage distribution.

| Mechanism | Increased confluence | Cell proliferation | Asymmetric inheritance | Multistage internalisation | Carrying capacity | Cell cycle | Nanoparticle recycling |
| --- | --- | --- | --- | --- | --- | --- | --- |
| Effect | Enhanced dosage depletion | Reduced uptake, broader dosage distribution | Multimodal, broader dosage distribution | Maintains Poisson dosage distribution | Narrower dosage distribution | Narrower dosage distribution | Maintains Poisson dosage distribution |
| Analytic results | Yes | Yes | Yes | Yes | No | No | No |

Biological heterogeneity can be explicitly included in the model by allowing the rate parameters of biological processes to follow specified distributions [8, 9]. However, we leave this extension for future work, noting that the analytic results provide an efficient method for analysing the confounding effects of biological heterogeneity and heterogeneity arising from the stochastic nature of nanoparticle-cell interactions. While we have examined a broad suite of biological mechanisms, it would be instructive to further develop the mathematical model to incorporate other potentially relevant phenomena such as sedimentary nanoparticle transport [36], transyctosis and degradation [11], and cell death. Additionally, we could extend the model to include internalisation rates that vary as a function of, for example, cell confluency [29] or cell cycle stage [43]. However, in the absence of experimental data to guide our choice of the modified rates, we leave this for future work. Here, we have primarily focused on theoretical analyses of the model. However, there is significant potential for interfacing the model with experimental data to draw conclusions about the sources of heterogeneity present. Such investigations will provide quantitative estimates of biological parameters in the case where the experimental data is sufficient to parameterise the appropriate model. Moreover, in the case where the data is not sufficient, we can employ the model to guide experimental protocol with respect to the type and frequency of data that is necessary. This will require considerable experimental effort, and is well beyond the scope of this manuscript, but remains an interesting avenue for future research.

## Code availability

All code to perform the model simulations is available on GitHub at https://github.com/celia2dow/Nanobio_Research_Project.

## Acknowledgements

The authors would like to acknowledge the generous guidance, mentoring and support provided by the late Prof. Edmund Crampin. Edmund taught us to think deeply about biological problems, with a rigour informed by his background in physics. He is terribly missed, and we hope to honour his legacy by continuing to apply his lessons to help unravel new problems.

The authors thank Prof. Sharon Lewin for the provision of laboratory facilities for P.M.C. The authors thank the anonymous referees for their helpful feedback.

## Appendix A. Experimental details

For the data in Figure 1, we aimed to assess the internalisation of 100 nm nanoparticles by Jurkat T cells as a representative example of quantitative nanoparticle-cell interaction data. To this end, 5 *μ*g 100 nm Absolute Mag™ Protein G Magnetic Particles (Creative Diagnostics, WHM-X035) were functionalised with 450 ng of AlexaFluor™ 647-labelled mouse-anti-human Transferrin Receptor (CD71) antibody (566724, BD Biosciences) in 10 mM phosphate buffer, pH 7.4 supplemented with 1 mg/mL bovine serum albumin for 30 minutes, then washed to remove excess antibody using magnetic separation and sonicated briefly. CD71 is well-described as a surface receptor that undergoes extensive receptor-mediated endocytosis [59], and as such can be used to trigger nanoparticle uptake into Jurkat T cells. In a 96-well plate format, 2.5 × 10^5^ Jurkat T cells were plated per well at a concentration of 1.25 × 10^6^ cells/mL, then incubated with CD71-functionalised nanoparticles at a cell : nanoparticle ratio of 1 : 100 for 1, 2, 4, 8, 24 or 48 hours. Cells were subsequently kept on ice to inhibit further nanoparticle internalisation, after which unbound nanoparticles were washed off and cells were stained with a fluorescent, phycoerythrin (PE)-labelled goat-anti-mouse antibody (P-852, Thermofisher Scientific). This secondary antibody co-labels any CD71-functionalized nanoparticles that have bound to the cell surface but have not been internalised. Fluorescent signals of both AlexaFluor™ 647 and PE were subsequently analysed using flow cytometry as a measure of associated and surface-bound nanoparticles, respectively, and processed via an arcsinh transform. No gating was applied during further analysis.

The data used in Figure 11 has been published previously and the experimental protocol and details can be found in the paper by Faria *et al*. [24].

## Appendix B. Corresponding differential equation models

The mean behaviour in our simulation model is consistent with the behaviour in various previously-proposed differential equation models of nanoparticle-cell interactions. Here we detail the conditions under which the previously-proposed models are equivalent to the mean behaviour in our simulation model. The key difference, however, is that our modelling framework is stochastic and allows us to analyse the expected dosage distributions, which cannot be obtained from deterministic differential equation models.

The differential equation model that is equivalent to the mean behaviour in the most general form of our simulation framework is

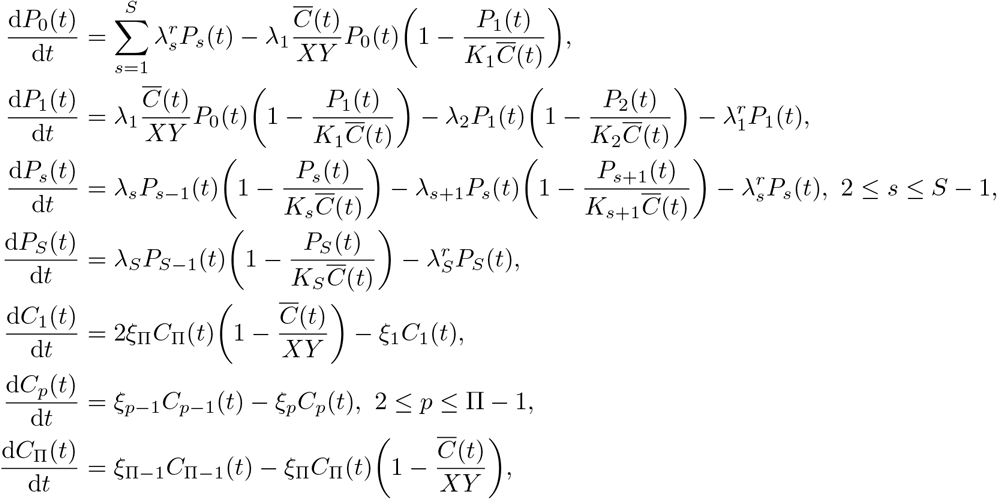

with initial conditions

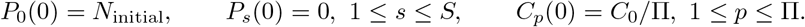

where 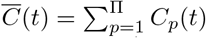 is the total number of cells, *C_p_*(*t*) is the number of cells in the *p*th stage of the cell cycle, *ξ_p_* = *P_p_/τ*, and *P_s_*(*t*) is the total number of nanoparticles in the *s*th stage of internalisation. Note that this is different to the number of nanoparticles per cell, but this can easily be recovered by 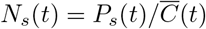. All other parameters are as defined in main manuscript.

While this system of equations may appear complicated, it is consistent with a number of previously-proposed models of nanoparticle-cell interactions. For example, the classic model of Wilhelm *et al*. [28] can be recovered under certain simplifying assumptions. Specifically, if *N*_initial_ is sufficiently large such that *P*_0_(*t*) ≈ *N*_initial_, *ξ_p_* = 0 ∀_*p*_, *S* = 2, and 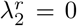. This is equivalent to assuming that the dosage available to the cells is approximately constant with time (i.e. no dosage depletion), there is no cell proliferation in the experiment, nanoparticle internalisation can be represented via a two-step process, and that there is no recycling of nanoparticles once in the final stage of internalisation. Under these assumptions, the general form of our model becomes

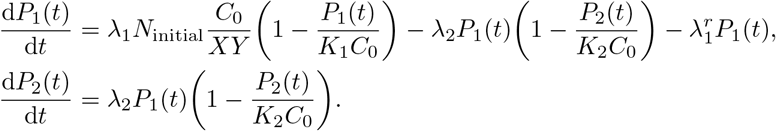

Once converted from total nanoparticles to nanoparticles per cell (i.e. dividing through by *C*_0_) and collecting parameters together, this reduced model is equivalent to the experimentally-validated model presented by Wilhelm *et al*. [28]. In Table 2 we provide a list of models that the mean behaviour of our model is equivalent to under various simplifying assumptions, with information about which assumptions are made and whether the model has been validated against experimental data.

**Table 2:**
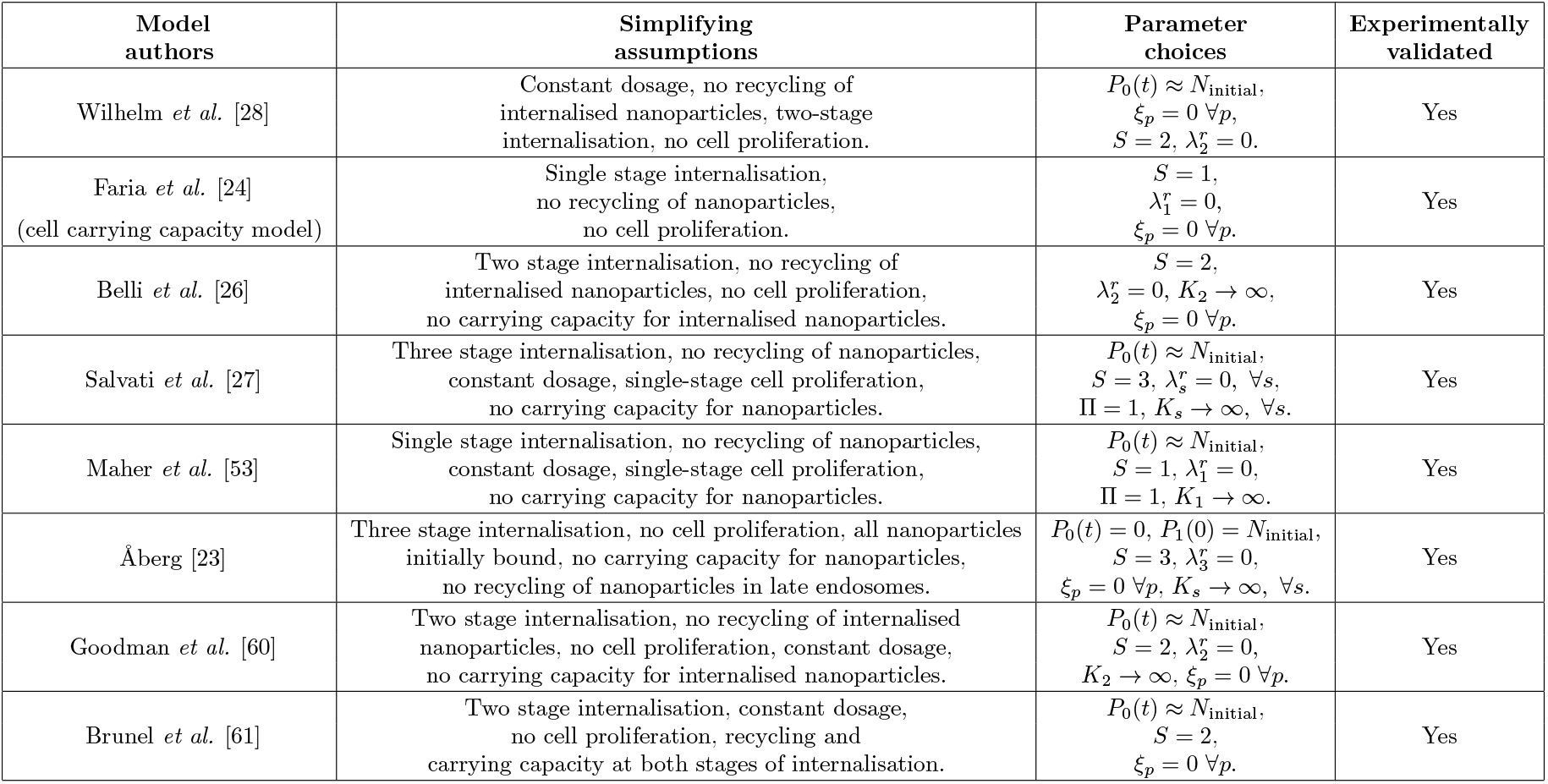
Details of the assumptions under which the mean behaviour in our simulation framework is equivalent to previously-published models of nanoparticle-cell interactions, and whether or not the previously-published model is validated against experimental data.

## Appendix C. Choice of the number of cell proliferation stages

Here we present results that highlight the impact of the choice of the number of stages in the cell proliferation model. In particular, in Figure 13, we examine the fraction of cells that have experienced a specific number division events at each time for three different choices of Π. We consider a single stage proliferation model (Π = 1), a model that represents the four stages of the cell cycle (Π = 4), and the model with Π = 24 considered previously in the manuscript. An increase in the number of stages in the proliferation process corresponds to a more consistent number of divisions experienced across the population. As noted based on the results presented in Figure 4, this corresponds to a decrease in the width of the dosage distribution, relative to a single stage model of proliferation. This suggests that the analytic results, which require an assumption of exponential waiting times between division events, will provide a slight overestimation of the expected heterogeneity.

**Figure 13:**
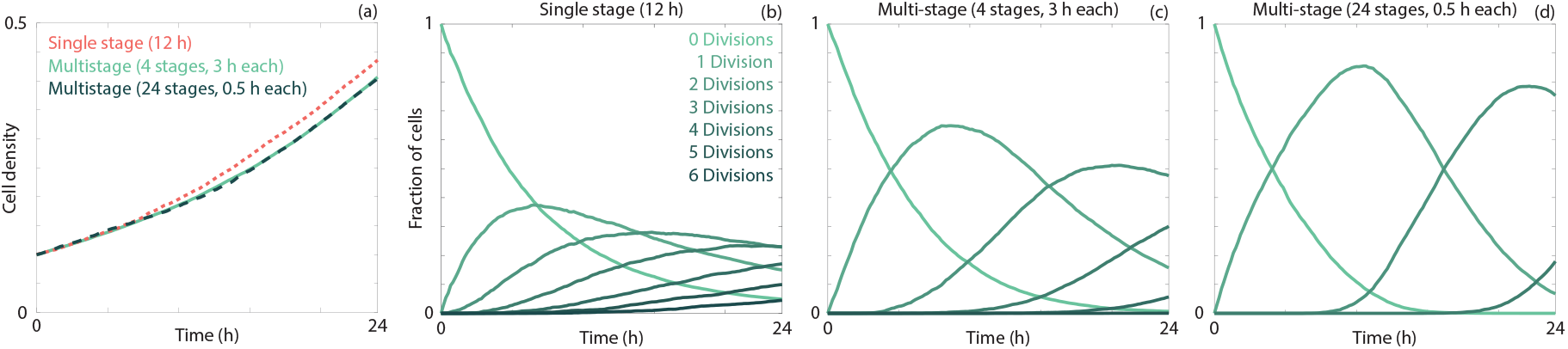
Comparison of different number of stages in the cell proliferation model. (a) Evolution of the cell density (i.e. number of cells divided by number of lattice sites) over time for the single stage model (red), the multistage model with Π = 4 (light green), and the multistage model with Π = 24 (dark green), such that the average time between cell divisions is constant. (b)-(d) The fraction of cells that have experienced 0, 1, 2, 3, 4, 5 or 6 division events over time for (b) the single stage model,(c) the multistage model with Π = 4, and (d) the multistage model with Π = 24.

